# Bacterial cell surface nanoenvironment requires a specialized chaperone to activate a peptidoglycan biosynthetic enzyme

**DOI:** 10.1101/2023.10.06.561273

**Authors:** Thomas Delerue, Sylvia Chareyre, Vivek Anantharaman, Michael C. Gilmore, David L. Popham, Felipe Cava, L. Aravind, Kumaran S. Ramamurthi

**Affiliations:** Laboratory of Molecular Biology, National Cancer Institute, National Institutes of Health, Bethesda, Maryland, USA; National Center for Biotechnology Information, National Library of Medicine, National Institutes of Health, Bethesda, Maryland, USA; The Laboratory for Molecular Infection Medicine Sweden (MIMS), Umeå Center for Microbial Research (UCMR), Science for Life Laboratory (SciLifeLab), Department of Molecular Biology, Umeå University, Umeå, Sweden; Department of Biological Sciences, Virginia Tech, Blacksburg, Virginia, USA

**Keywords:** SpoIVA, SpoVM, DivIVA, MreB, FtsZ, Clostridium

## Abstract

*Bacillus subtilis* spores are produced inside the cytosol of a mother cell. Spore surface assembly requires the SpoVK protein in the mother cell, but its function is unknown. Here, we report that SpoVK is a dedicated chaperone from a distinct higher-order clade of AAA+ ATPases that activates the peptidoglycan glycosyltransferase MurG during sporulation, even though MurG does not normally require activation by a chaperone during vegetative growth. MurG redeploys to the spore surface during sporulation, where we show that the local pH is reduced and propose that this change in cytosolic nanoenvironment necessitates a specific chaperone for proper MurG function. Further, we show that SpoVK participates in a developmental checkpoint in which improper spore surface assembly inactivates SpoVK, which leads to sporulation arrest. The AAA+ ATPase clade containing SpoVK includes other dedicated chaperones involved in secretion, cell-envelope biosynthesis, and carbohydrate metabolism, suggesting that such fine-tuning might be a widespread feature of different subcellular nanoenvironments.

## INTRODUCTION

The use of cytosolic molecular chaperones in aiding proteins that fold improperly in response to harsh environmental conditions has been extensively studied in the context of bacterial stress responses (2–4). Insults such as oxidative stress and extreme temperature and pH can cause protein denaturation and, ultimately, loss of function. As a defense, bacteria have evolved specialized proteins, termed chaperones, that aid protein folding, preventing the aggregation of misfolded proteins, and even resolving protein aggregates. An important group of these chaperones belongs to the AAA+ family, which includes those that assist protein folding in mitochondria, the endoplasmic reticulum, and bacterial cells (5–7).

Spore formation is a developmental program that certain Gram-positive bacteria initiate when faced with starvation (8–10). In *Bacillus subtilis*, sporulation initiates with an asymmetric division event that divides the progenitor cell into two unequally sized daughter cells that will display different cell fates: a smaller forespore that will mature into a dormant cell type called the “spore” and a larger mother cell that will eventually lyse (Fig. 1A). Next, the mother cell engulfs the forespore and builds two concentric shells around the forespore that will eventually protect the mature spore from the environment: an inner peptidoglycan “cortex” built between the double membrane envelope surrounding the forespore, and an outer proteinaceous “coat” built around the outer forespore membrane. The coat is a complex structure consisting of ∼80 mother cell-produced proteins (11) whose basement layer is built using SpoIVA, a cytoskeletal protein (12–14) that hydrolyzes ATP to irreversibly assemble around the developing forespore (15–17). Eventually, the mother cell lyses in a programmed manner and releases the mature, dormant spore.

**Figure 1.**
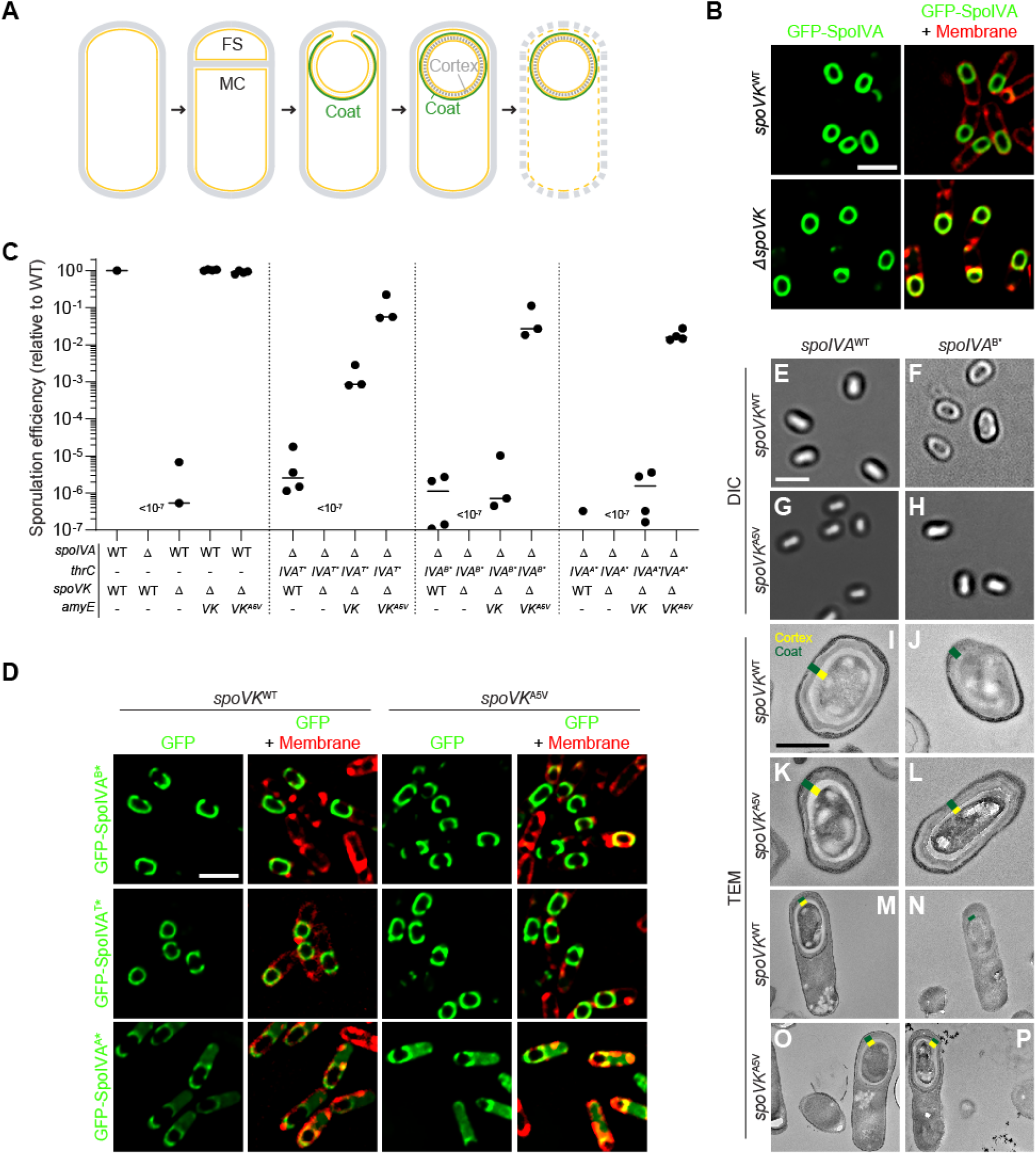
Point mutant in *spoVK* suppresses spore envelope assembly defects caused by mis-assembly of the spore coat basement layer. (A) Schematic representation of sporulation in *Bacillus subtilis*. Asymmetric division results in the formation of a small forespore (FS) and larger mother cell (MC). A proteinaceous shell, the “coat” (green) is first constructed on the outer forespore membrane; a peptidoglycan shell, the “cortex” (gray dashes) is later constructed between the two membranes surrounding the forespore. The mother cell ultimately lyses, releasing the mature forespore into the environment. Membranes are depicted in yellow; peptidoglycan cell wall is depicted in gray. (B) Subcellular localization of GFP-SpoIVA (green) in presence (top panels) or absence (bottom panels) of *spoVK*, 3.5 h after induction of sporulation (strains: KR160 and TD549). Membranes visualized using fluorescent dye FM4-64 (red; right panels). (C) Sporulation efficiencies, determined as resistance to heat, relative to WT (strain PY79). Strain genotypes at *spoIVA* and *spoVK* loci are indicated below the graph; *thrC* and *amyE* are ectopic chromosomal loci used to complement *spoIVA* and *spoVK* deletions, respectively, with different alleles of those genes. Bars represent mean values. Strains: PY79, KP73, TD520, TD513, TD514, JPC221, TD524, TD530, TD531, TD523, TD528, TD529, KR438, TD817, TD818, and TD819. (D) Subcellular localization of GFP-SpoIVA^B*^, GFP-SpoIVA^T*^, or GFP-SpoIVA^A*^ variants that fails to polymerize, in cells producing SpoVK^WT^ (left) or SpoVK^A5V^ (right; strains: TD845, TD846, TD854, TD848, TD849, and TD855) 4 h after induction of sporulation. Left: fluorescence from GFP; right: overlay, fluorescence from GFP and membranes visualized with FM4-64. Size bars: 2 µm. (E-L) Released spores visualized using (E-H) differential interference contrast (DIC) light microscopy (size bar: 2 µm), or (I-L) transmission electron microscopy (TEM; size bar: 500 nm) harboring WT (left) or B* alleles (right) of *spoIVA* in the presence of *spoVK^WT^* (E-F; I-J) or *spoVK^A5V^* (G-H; K-L). (M-P) TEM images of strains in (I-L) at 5.5 h after induction of sporulation. Coat (green) and cortex (yellow) are marked, when present, in (I-P). Strains: PY79, JB103, TD514, and TD529. Strain genotypes are listed in Table S2.

Although the final structure of the cortex peptidoglycan is different from vegetative peptidoglycan, it is built using the same precursors (18). During vegetative growth, the final precursor, the membrane-bound lipid II, is synthesized in the cytosol and flipped to the oxidizing environment of the cell surface; during sporulation, lipid II is synthesized in the mother cell and flipped into the forespore intermembrane space. Vegetative growth requires peptidoglycan insertion in two different locations: at the lateral edge of the growing cell and at mid-cell during cell division (19–21). Accordingly, specific transpeptidases and transglycosylases are tasked with attaching new cell wall material into the appropriate location and must coordinate their activity with cell membrane formation or the cell division machinery. Although a link between coat assembly and the transglycosylases and transpeptidases that mediate cortex assembly has not been formally established, cortex assembly is nonetheless subject to the coat assembly checkpoint. In this checkpoint, cortex assembly will not initiate unless the basement layer of the coat, composed of SpoIVA, properly polymerizes (22–24). Recently, we reported that the SpoIVA-associated protein SpoVID (25–27) physically monitors the polymerization state of SpoIVA via the C-terminus of SpoVID, which curiously harbors a peptidoglycan-binding domain called LysM (28). When SpoIVA polymerizes properly, the C-terminus of SpoVID binds to the polymerized SpoIVA, occluding the LysM domain. However, if SpoIVA mis-assembles, the C-terminus of SpoVID is liberated, thereby exposing the LysM domain which binds to and sequesters the lipid II precursor in the mother cell cytosol, thereby blocking cortex assembly. Although SpoVID molecules (and therefore LysM domains) vastly outnumber the estimated number of lipid II molecules at any given time (28), it was not clear how the checkpoint could remain functional in the face of additional lipid II synthesis that may overwhelm the lipid II sequestration capacity of SpoVID.

In this study, we employed a genetic strategy to identify additional factors that participate in the coat assembly checkpoint, which yielded the previously uncharacterized AAA+ chaperone SpoVK (29). We report that SpoVK activates the lipid II synthase MurG. During sporulation, MurG redeploys from the mother cell plasma membrane to the forespore surface (30) to generate lipid II for cortex assembly, but MurG does not normally require an activating chaperone during vegetative growth. Examining the pH of the mother cell cytosol immediately adjacent to the forespore surface revealed a cytosolic nanoenvironment where the pH was lower than the rest of the mother cell cytosol. We propose that MurG is largely nonfunctional in this nanoenvironment and requires activation by SpoVK to function during sporulation. Thus, the contiguous cytosol of the mother cell during sporulation is not uniform and the presence of certain cytosolic nanoenvironments may necessitate assisted protein folding for select proteins. Further, we show that this requirement for SpoVK at the forespore surface offers an additional level of regulation wherein the coat assembly checkpoint can inactivate SpoVK upon sensing coat assembly mistakes to prevent the accumulation of lipid II.

## RESULTS

### Suppressor mutation in spoVK restores cortex assembly caused by defective SpoIVA variants

SpoIVA hydrolyzes ATP to polymerize irreversibly on the forespore surface to form a stable platform upon which the spore coat assembles (14, 16). Since successful initiation of coat assembly is required to trigger cortex assembly, strains of *B. subtilis* harboring defective SpoIVA variants that do not polymerize properly not only fail to assemble the spore coat, but also do not construct a cortex and are therefore unable to sporulate (31). Previously, we exploited this phenotype to isolate a spontaneous suppressor that would correct the sporulation defect of SpoIVA^T*^, a defective variant with a disrupted “sensor threonine” that binds, but does not hydrolyze, ATP (17). This genetic selection yielded a loss-of-function mutation in the *spoVID* gene (25). We previously presented evidence that SpoVID participates in a checkpoint that monitors coat assembly and, upon sensing a coat assembly defect, would arrest cortex assembly by sequestering the peptidoglycan intermediate lipid II via a C-terminal LysM domain that is unmasked only when the coat mis-assembles (28). To identify additional factors involved in the communication between coat assembly and cortex formation, we used a similar genetic strategy to isolate additional suppressor mutations that would correct the sporulation defect of *spoIVA^T*^*, but this time we used a strain harboring two copies of *spoVID* to avoid isolating loss-of-function mutations in *spoVID*. To isolate suppressors, we grew cells in sporulation media and removed nonsporulating and poorly sporulating cells by exposure to 80 °C for 20 min. Surviving cells were then enriched by repeated dilution of the heat-killed culture into fresh sporulation media, where survivors could germinate and re-sporulate. Whole genome sequencing revealed an extragenic mutation in the *spoVK* gene wherein a cytidyl to thymidyl transition changed the specificity of codon 5 from alanine to valine.

SpoVK (previously misnamed SpoVJ) is a poorly studied sporulation protein produced in the mother cell under the control of the σ^E^ promoter; deletion of *spoVK* resulted in a severe sporulation defect, but its function has remained mysterious (29, 32). Later, SpoVK was classified as a member of the AAA+ family of P-loop NTPases (7). Since SpoVK had not been previously implicated in coat assembly, we first examined the subcellular localization of the coat protein SpoIVA in the presence or absence of SpoVK. GFP-SpoIVA localizes to the forespore surface (Fig. 1B). In the absence of SpoVK, GFP-SpoIVA localization was similar to WT, indicating that SpoVK is not involved directly in coat basement layer assembly. Nonetheless, deletion of *spoVK* resulted in a ∼10^6^-fold defect in sporulation efficiency relative to WT (Fig. 1C, lane 3), even though the basement layer appeared to assemble normally.

Next, we examined the allele specificity of suppression by *spoVK^A5V^*. In the presence of WT SpoIVA, complementation of the *spoVK* deletion *in trans* with a single copy of either *spoVK or spoVK^A5V^* at an ectopic chromosomal locus (*amyE*) restored sporulation efficiency to a level similar to WT (Fig. 1C, lanes 4-5). In the presence of *spoIVA^T^*\* (the *spoIVA* allele used to isolate the *spoVK^A5V^* suppressor mutation) or alleles of *spoIVA* in which the Walker B or Walker A motifs were disrupted (*spoIVA^B^*\* and *spoIVA^A^*\*), which abrogates ATP hydrolysis or binding, respectively (17, 33), introducing the *spoVK^A5V^* allele also suppressed the sporulation defect caused by the mutant *spoIVA* allele (Fig. 1C, lanes 6-17), indicating that suppression by the *spoVK^A5V^* mutation is not allele-specific.

SpoIVA variants that are defective in ATP hydrolysis or binding display subtle localization defects in vivo: GFP-SpoIVA^B*^ and GFP-SpoIVA^T*^ fail to fully encircle the forespore and GFP-SpoIVA^A*^ is predominantly cytosolic (Fig. 1D). Introducing SpoVK^A5V^ did not restore proper localization of these SpoIVA variants (Fig. 1D), despite largely correcting the sporulation defect caused by these SpoIVA variants (Fig. 1C). When viewed by light microscopy, mature WT spores appear as “phase bright” (Fig. 1E) due to the exclusion of water from the spore core, which is maintained by a robust cortex that may be observed using transmission electron microscopy (Fig. 1I, 1M; indicated in yellow). In contrast, cells harboring a defective *spoIVA* allele, such as *spoIVA^B^*\*, that mis-assemble the coat produce “phase gray” spores (Fig. 1F) and fail to build a cortex (Fig. 1J, N). The presence of SpoVK^A5V^ did not affect the morphology of otherwise WT cells (Fig. 1G, 1K, 1O), but did restore phase brightness and cortex assembly to cells that harbored *spoIVA^B^*\* (Fig. 1H, 1L, 1P). The results are consistent with a model in which SpoVK participates in the pathway linking coat and cortex assembly, and that *spoVK^A5V^* is a gain-of-function allele that permits the cell to build a functional cortex even if the basement layer of the spore coat is defective.

### SpoVK is an ephemeral forespore-associated protein involved with cortex assembly

In the absence of SpoVK, cells produced phase gray spores which, when viewed by TEM, did not harbor a cortex (Fig. 2A, top row). At 5.5 h after sporulation initiation, WT cells, viewed by TEM, produced a thick cortex, but cells lacking SpoVK instead displayed a thin layer of peptidoglycan surrounding the forespore (Fig. 2A, bottom row, arrow) that excluded the negative stain. To test if this thin peptidoglycan layer was chemically different from cortex peptidoglycan, we extracted cortex peptidoglycan from sporulating WT and Δ*spoVK* cells, digested with mutanolysin, identified and quantified the muropeptides by their characteristic elution times when separated by HPLC, and calculated peptidoglycan structural parameters (34). The levels of three muropeptides that are characteristic of cortex peptidoglycan (muramic acid present as lactam, disaccharide units in tetrasaccharide, and disaccharide units in hexasaccharide) were reduced in the Δ*spoVK* cells compared to WT (Fig. 2B, Table S1). In contrast, levels of three characteristics that are underrepresented in cortex peptidoglycan but are present in vegetative peptidoglycan and in the innermost “germ cell wall” layer of the cortex (muramic acid with tetrapeptide, disaccharide units in disaccharide, and peptide present as tripeptide) were higher in the Δ*spoVK* cells compared to WT (Fig. 2B). The results suggest that SpoVK is required for producing the structurally distinct cortex peptidoglycan during sporulation.

**Figure 2.**
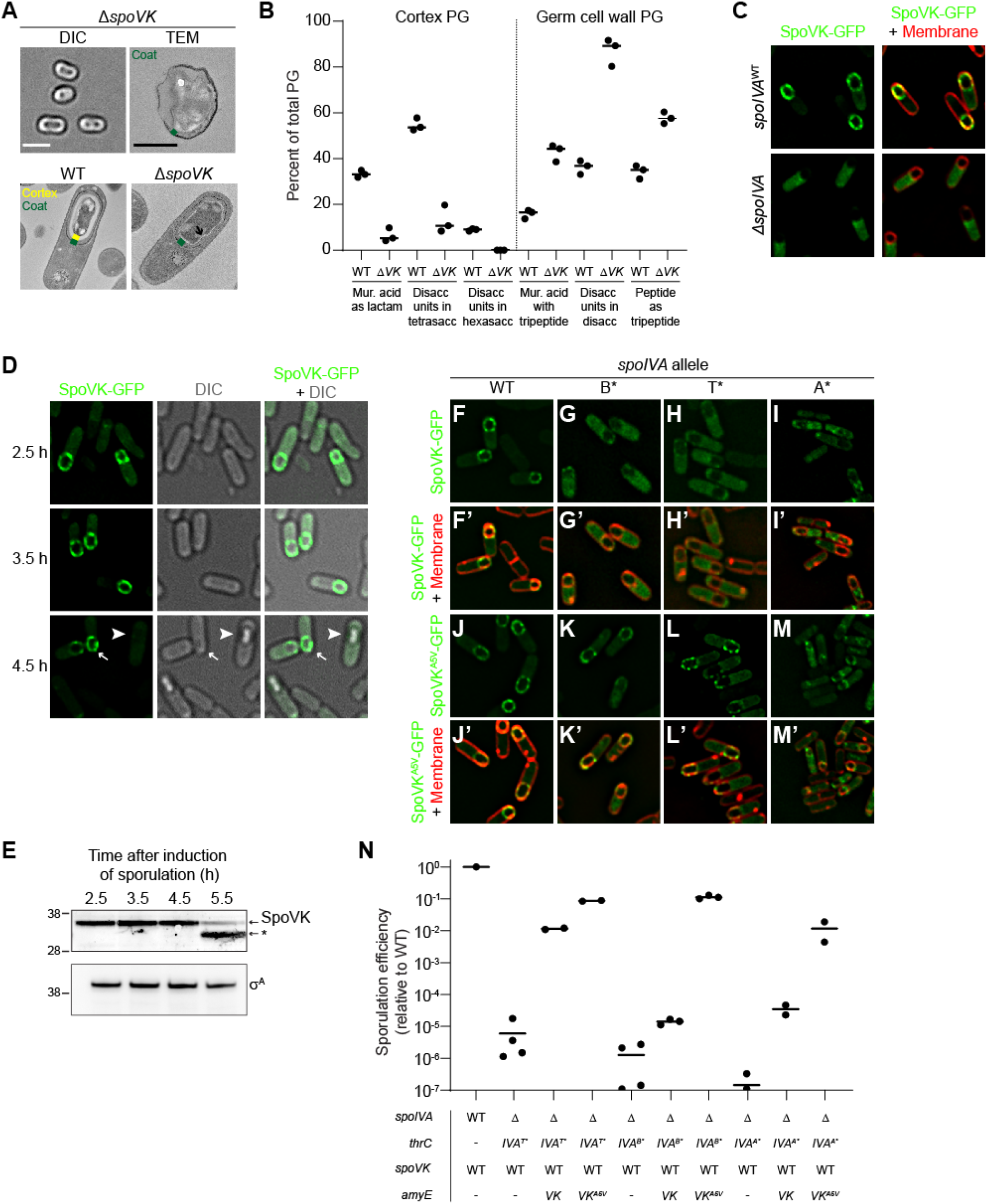
SpoVK is an ephemeral sporulation protein. (A) DIC light microscopy (left; size bar: 2 µm) or TEM images (F; size bar: 500 nm) of released spores (top panels) containing deletion of *spoVK* (strain TD520). Bottom panels: TEM images of WT (left) or *spoVK* deletion strain at 5.5 h after induction of sporulation. Green in the TEM images indicates coat; yellow indicates cortex. (B) Structural parameters of spore peptidoglycan produced by WT or Δ*spoVK* (strains PY79 and TD520, respectively). Peptidoglycan from developing spores was extracted 5 h after induction of sporulation, digested with mutanolysin, and separated by HPLC. Peaks of select muropeptides that are characteristic cortex or germ cell wall were integrated and depicted as a percent of total peptidoglycan (see Table S1 for complete analysis). Bars represent mean; data points represent an independent culture. (C) Subcellular localization of SpoVK-GFP in presence (top panels) or absence (bottom panels) of *spoIVA* 3.5 h after induction of sporulation (strains TD604 and TD652). Left: fluorescence from SpoVK-GFP; right: overlay of GFP fluorescence (green) and FM4-64 (red). (D) Subcellular localization of SpoVK-GFP at the indicated (left) time points after the induction of sporulation. Left panels: fluorescence from SpoVK-GFP; center: DIC; right: overlay of fluorescence and DIC. Arrowhead indicates a phase bright forespore; arrow indicates phase gray forespore (strain TD604). (E) Immunoblot of cell extracts of sporulating wild type *B. subtilis* (strain PY79) using anti-SpoVK or anti-σ^A^ antisera, from cells harvested 2.5 h, 3.5 h, 4.5 h, and 5.5 h. Relative mobility of molecular weight markers (kDa) indicated to the let; asterisk indicates a likely degradation product of SpoVK. (F-M’) Subcellular localization of (F-I’) SpoVK-GFP or (J-M’) SpoVK^A5V^-GFP in the presence of (F-F’; J-J’) SpoIVA^WT^ or (G-M’) SpoIVA variants that fails to polymerize (IVA^B*^, IVA^T*^ or -IVA^A*^) 3.5 h after the induction of sporulation (strains: TD675, TD682, TD684, and TD836). (F-M) GFP fluorescence; (F’-M’) overlay, GFP fluorescence and FM4-64. (N) Sporulation efficiencies, determined as resistance to heat, relative to WT (PY79). Strain genotypes at *spoIVA* and *spoVK* loci are indicated below the graph; *thrC* and *amyE* are ectopic chromosomal loci used to complement *spoIVA* and *spoVK* deletions, respectively, with different alleles of those genes. Bars represent mean values; data points represent an independent culture (Strains PY79, JPC221, TD563, TD564, JPC75, TD557, TD558, KR438, TD859, and TD860).

We next examined the subcellular localization of SpoVK-GFP. In otherwise WT cells, SpoVK-GFP localized as puncta at the periphery of the forespore (Fig. 2C). In the absence of SpoIVA, SpoVK-GFP was instead localized in the mother cell cytosol (Fig. 2C). This genetic dependence on SpoIVA for forespore localization is characteristic of spore coat proteins (11). To test if, like other coat proteins, SpoVK remains associated with the mature spore, we examined SpoVK-GFP localization in sporulating cells at different time points. At 2.5 h and 3.5 h after induction of sporulation, SpoVK-GFP remained associated with the forespore periphery (Fig. 2D). However, at later time points, specifically in cells that had elaborated a phase-bright forespore, SpoVK-GFP was undetectable (Fig. 2D). To ensure that the disappearance of SpoVK-GFP in these cells was not due to loss of fluorescence of the GFP fusion, we examined the presence of native SpoVK in sporulating cells by immunoblotting. SpoVK was detectable in cell extracts at early time points, but at 5.5 h after induction of sporulation, the level of full length SpoVK was diminished and a faster migrating species, likely a SpoVK degradation product, accumulated (Fig. 2E). Thus, SpoVK temporarily associates with the forespore surface in a SpoIVA-dependent manner and is likely degraded once cells elaborate a mature cortex.

To understand the role of the A5V substitution in suppressing defective alleles of *spoIVA*, we examined the subcellular localization of SpoVK^A5V^-GFP. In the presence of ATP hydrolysis-defective variants of SpoIVA (B* and T*), SpoVK-GFP mis-localized in the mother cell cytosol (Fig. 2F-H, F’-H’), but SpoVK^A5V^-GFP largely correctly localized to the forespore surface despite a defective coat basement layer (Fig. 2J-L, 2J’-2L’). In the presence of the ATP binding-defective SpoIVA variant (A*), the mis-localization of SpoVK-GFP was more pronounced (Fig. 2I, 2I’), but the A5V substitution nonetheless restored at least partial localization to the forespore periphery (Fig. 2M-M’). We next tested if the A5V substitution resulted in a gain of function in SpoVK by examining sporulation efficiency in merodiploid strains. In the presence of SpoIVA variants harboring defects in the nucleotide-binding pocket, addition of a single copy of WT *spoVK* improved the sporulation efficiency slightly (*spoIVA^B^*\*) or ∼10^2^-10^3^-fold (*spoIVA^A*^* and *spoIVA^T^*\*); the addition of *spoVK^A5V^* improved sporulation efficiency even further (∼10^4^-10^5^ fold; Fig. 2N). We therefore conclude that SpoVK is a short-lived spore coat-associated protein and that *spoVK^A5V^* is a gain-of-function allele.

### SpoVK is a AAA+ ATPase specific to sporulating Firmicutes

While SpoVK has been recognized as an AAA+ ATPase, its affinities within this vast and functionally diverse family of P-loop NTPase enzymes remain poorly understood. Hence, we conducted a systematic evolutionary and structural investigation of SpoVK using sensitive sequence profile analysis, phylogenetic tree construction, and structural modeling. Profile-profile comparisons revealed that it is a member of the “Classical AAA’’ assemblage prototyped by the proteasomal subunits, the chaperone CDC48 and FtsH (HHpred p=97-99.5%). Phylogenetic analysis revealed that within that assemblage, SpoVK belongs to a higher-order clade that specifically unites it with SpoVK-like ATPases, the ribulose bisphosphate carboxylase (RuBisCO) chaperone CbbX and its relatives, the Type VII secretion system (T7SS) EccA-like chaperones, and other poorly characterized eukaryotic proteins predicted to play a role in RNA processing (Fig. 3A). This higher-order clade is, in turn, a sister group of the FtsH AAA+ domains within the Classical AAA assemblage, which share the unifying structural feature of a bihelical hairpin immediately upstream of the Walker A motif (Fig. S1A-B). The apex of this hairpin binds the adenine moiety of the bound nucleotide. This clade is unified by and distinguished from FtsH by: (i) a conserved tyrosine in the sensor-1 region as part of a GY signature; (ii) a conserved asparagine 4 residues upstream of the first arginine finger; and (iii) the presence of the second arginine finger (sensor-2) that is usually lost in FtsH (Fig. S1B). Within the higher-order clade, SpoVK and related proteins emerged as most closely related to the chaperone CbbX (Fig. 3A, S1C).

**Figure 3.**
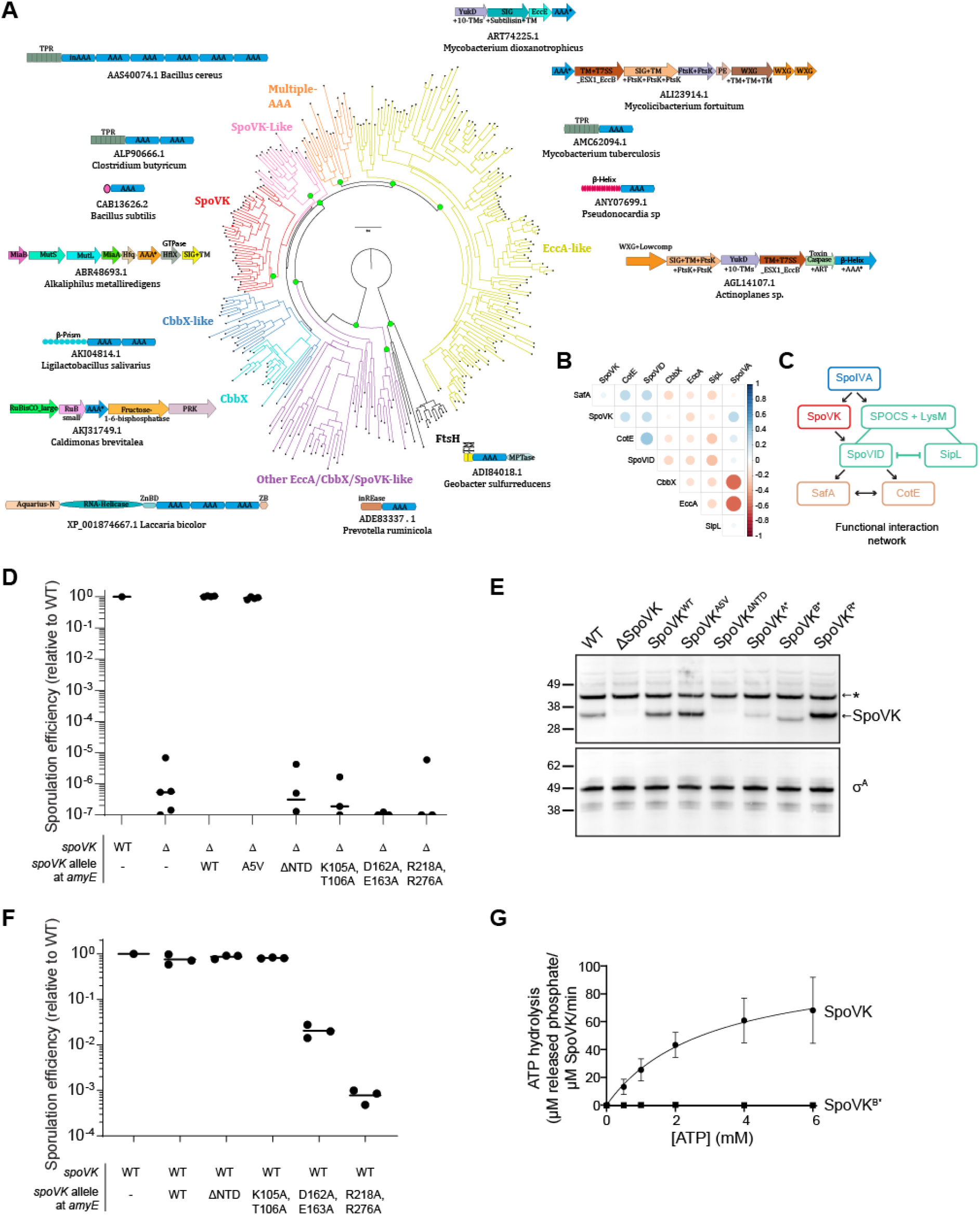
SpoVK is a functional triple AAA+ protein. (A) Phyletic analysis of the SpoVK-EccA-like clade. A phylogenetic tree of a representative set of AAA+ domains in the SpoVK-EccA-like clade is shown. The tree shows the relationship between the SpoVK, SpoVK-like, CbbX, CbbX-like, and EccA-like families and the closest outgroup FtsH. Branches are colored according to the families. Select operonic arrangements or domain architectures of proteins are shown and placed near the branch of the tree in which they or their orthologs occur. Each arrow in the operon is a gene coding for the protein. The accession and organism of the AAA+ containing protein in the operon, marked with an asterisk, is shown below each operon or domain architecture. ZnBd: zinc-binding. (B) Correlation matrix of SpoIVA, SpoVK, CbbX, EccA, SafA, and SPOCS domain-containing proteins SpoVID, CotE, and SipL/DUF3794 was computed from their phyletic pattern vectors. Color intensity and the size of the circle are proportional to the correlation coefficient. (C) A correlation network of SpoIVA, SpoVK, CbbX, EccA, SafA, and SPOCS domain-containing proteins SpoVID, CotE, and SipL/DUF3794 was computed from this matrix and the phyletic pattern vectors. The nodes and edges are colored as per their functional subgroup in the sporulation system. The arrowheads indicate positive correlations and point from the phyletically more widespread to the less widespread proteins. The flat heads indicate negative correlations. Lines indicate a split in a functionally equivalent family into sub-groups (e.g., SipL and SpoVID). (D) Sporulation efficiencies, determined as resistance to heat, relative to WT (PY79). Strain genotypes at the *spoVK* locus are indicated below the graph; *amyE* is an ectopic chromosomal locus used to complement the *spoVK* deletion strain with the indicated allele of *spoVK*. Bars represent mean values. Strains: PY79, TD520, TD513, TD514, TD574, TD575, TD576, and TD578. (E) Immunoblot of cell extracts of sporulating *B. subtilis* using anti-SpoVK or anti-σ^A^ antisera, from cells harvested at 4 h after induction of sporulation. Strains: PY79, TD520, TD513, TD514, TD574, TD575, TD576, and TD578. (F) Sporulation efficiencies, determined as resistance to heat, relative to WT (PY79). Strain genotypes at *spoVK* locus are indicated below the graph; *amyE* is an ectopic chromosomal locus used to produce a merodiploid strain containing two alleles of *spoVK*. Bars represent mean values. Strains: PY79, TD597, TD598, TD599, TD600, and TD602. (G) Saturation curve for ATP hydrolysis by purified SpoVK and SpoVK variant harboring a disrupted Walker B motif (D162A, E163A). Purified SpoVK or SpoVK^D162A, E163A^ was incubated with increasing concentrations of ATP, and nucleotide hydrolysis was assayed by measuring the generation of free phosphate. Data were fit to the Michaelis-Menten enzyme saturation model.

The N-terminal region of this clade of AAA+ proteins shows considerable variability and features multiple alternative domains (Fig. 3A; e.g., TPRs and a β-helix in EccA, a β-prism in certain CbbX-like versions, and SF-1 RNA helicase domain in eukaryotic representatives) that are indicative of mediating interactions which might recruit the substrate. Our analysis revealed the SpoVK proteins have a short, unique N-terminal domain with 3 β-strands and an α-helix (Fig. S1B). The A5V substitution lies in this region and is predicted to stabilize the first strand of the N-terminal domain keeping with its gain-of-function phenotype. Our phylogenetic analysis also helped objectively discriminate SpoVK from other members of this clade like CbbX or EccA and thereby allowed us to accurately infer its phyletic patterns (Fig. S2). We used this revised phyletic pattern of SpoVK along with those of other related AAA+ ATPases and sporulation proteins to construct a phyletic correlation matrix and extract a functional interaction network from it. This indicated that SpoVK is only found in sporulating firmicutes and is part of a functional interaction network that includes SpoIVA, SpoVID, CotE, and SafA (Fig. 3B). The phyletic patterns further suggest that SpoVK might function downstream of SpoIVA and in parallel with SpoVID (or another factor that harbors a SPOCS and LysM domain such as SipL) in a sporulation specific network (Fig. 3C, Fig. S2). In contrast to the proteasomal, FtsH and CDC48 AAA+ ATPases with rather generic targets, the characterized members of the higher-order clade containing SpoVK function as chaperones with “narrow” roles directed at a dedicated target (like RuBisCo in the case of CbbX) or in a specific sub-cellular context (like EccA, which operates on T7SS substrates) (35). Hence, we predict that SpoVK might likewise function as a dedicated chaperone in a specific sporulation-related context.

### SpoVK displays ATPase activity *in vitro*

To test if SpoVK is a functional AAA+ NTPase, we first disrupted the conserved Walker A and Walker B motifs and the first arginine finger upstream of strand-5 (Fig. S1A). Disruption of any of these motifs resulted in severe sporulation defects, similar to deletion of the *spoVK* gene (Fig. 3D, lanes 6-8). Examination of cell extracts of sporulating cells by immunoblotting revealed that disruption of the Walker A or Walker B motifs, but not the arginine finger, reduced, but did not eliminate, accumulation of SpoVK protein relative to WT SpoVK (Fig. 3E, lanes 6-8). Next, we examined the role of the N-terminal domain (NTD; residues 2-41), which might recruit substrates as in other AAA+ proteins. Deletion of the NTD completely abrogated accumulation of SpoVK (Fig. 3E, lane 5), and consequently resulted in reduced sporulation efficiency, similar to deletion of the *spoVK* gene (Fig. 3D, lane 5).

Alphafold2 multimer modeling indicated that SpoVK is likely to assume a homo-hexameric functional state comparable to its closest sister clades such as CbbX (Figure S1D). We therefore tested the dominance of the *spoVK* mutant alleles in merodiploid strains that also contained the WT copy of *spoVK*. Co-expressing *spoVK*^ΔNTD^ or *spoVK*^A*^ did not reduce sporulation efficiency in cells harboring a WT copy of *spoVK* (Fig. 3F, lanes 3-4), presumably because these variants did not accumulate to appreciable levels in the cell. However, co-expression of *spoVK*^B*^ or *spoVK*^R*^ with WT spoVK resulted in 100-1000 fold reduction in sporulation efficiency (Fig. 3F, lanes 5-6). The genetic dominance of these alleles suggested that SpoVK is capable of oligomerizing in vivo and that, similar to other AAA+ proteins, introduction of ATPase-defective subunits can poison the function of the oligomerized chaperone (5).

Finally, we examined if SpoVK could hydrolyze ATP in vitro. Incubation of purified SpoVK-His_6_ with increasing concentrations of ATP produced a saturation curve that revealed a substrate turnover rate (*k*_cat_) of 104 ± 18 µM released phosphate min^-1^ µM^-1^ SpoVK (Fig. 3G). In contrast, purified SpoVK^B*^ did not appreciably hydrolyze ATP at any of the concentrations tested. Thus, as predicted by the sequence-structure analysis, the *in vivo* and *in vitro* results suggest that SpoVK is a functional AAA+ ATPase.

### SpoVK interacts with SpoVID and MurG

To elucidate the role of SpoVK during sporulation, we immunoprecipitated FLAG-tagged SpoVK from sporulating *B. subtilis* cultures and identified co-purifying proteins. To trap associated proteins, we performed the immunoprecipitation with FLAG-tagged SpoVK harboring a disrupted Walker B motif (SpoVK^B*^-FLAG), a strategy that we have successfully used previously to stabilize interactions between chaperones and potential substrates (36). Immunoprecipitation of SpoVK^B*^-FLAG from extracts of cells 4 h after induction of sporulation revealed two species specifically in the eluate that were not present when the immunoprecipitation was performed with cells producing SpoVK^B*^-His_6_: a ∼36 kDa band and a ∼65 kDa band (Fig. 4A). Trypsin digestion of these bands followed by mass spectrometry revealed, as expected, that the ∼36 kDa band was SpoVK^B*^-FLAG. The ∼65 kDa band was identified as SpoVID, a factor that we recently identified as participating in a sporulation checkpoint that monitors spore coat assembly (28). Immunoblotting the fractions using anti-SpoVID antibodies confirmed that SpoVID specifically co-purified with SpoVK^B*^-FLAG and the gain-of-function SpoVK^B*, A5V^-FLAG, but not SpoVK^B*^-His_6_ (Fig. 4B). To test the relationship between SpoVID and SpoVK in vivo, we first examined the subcellular localization of SpoVK-GFP. In the absence of SpoVID, SpoVK-GFP mis-localized on the mother cell-distal side of the forespore instead of uniformly localizing around the entire forespore surface (Fig. 4C). However, this mis-localization of SpoVK did not require the peptidoglycan-binding capability of the SpoVID LysM domain since a single amino acid substitution disrupting this function of the LysM domain (SpoVID^T532G^) did not result in SpoVK mis-localization (Fig. 4C). Nonetheless, expression of the LysM domain of SpoVID alone (SpoVID^501-575^) was sufficient to restore proper localization of SpoVK-GFP (Fig. 4C), indicating that the C-terminal LysM domain of SpoVID, but not a functional LysM domain (namely, one capable of binding peptidoglycan precursors) *per se*, influences the subcellular localization of SpoVK. We next examined the stability of SpoVK by immunoblotting. In WT cells, SpoVK was detectable at a low level; deletion of *spoIVA*, which results in spore coat mis-assembly, resulted in even lower levels of SpoVK, which was restored upon complementation of *spoIVA* at an ectopic locus (Fig. 4D, lanes 1-3). Deletion of *spoVID*, though, increased the level of SpoVK; this increased level of SpoVK was reversed upon complementation of *spoVID* at an ectopic locus (Fig. 4D, lane 4-5). Expression of the LysM domain of SpoVID alone, as the only copy of SpoVID, was sufficient to maintain low, ∼WT levels SpoVK, but additional truncation of the LysM domain resulted in increased levels of SpoVK (Fig. 4D, lanes 6-7). We conclude that the LysM domain of SpoVID, which is unmasked when the spore coat fails to assemble properly (28), is simultaneously required for proper localization of SpoVK and negatively influences the stability of SpoVK.

**Figure 4.**
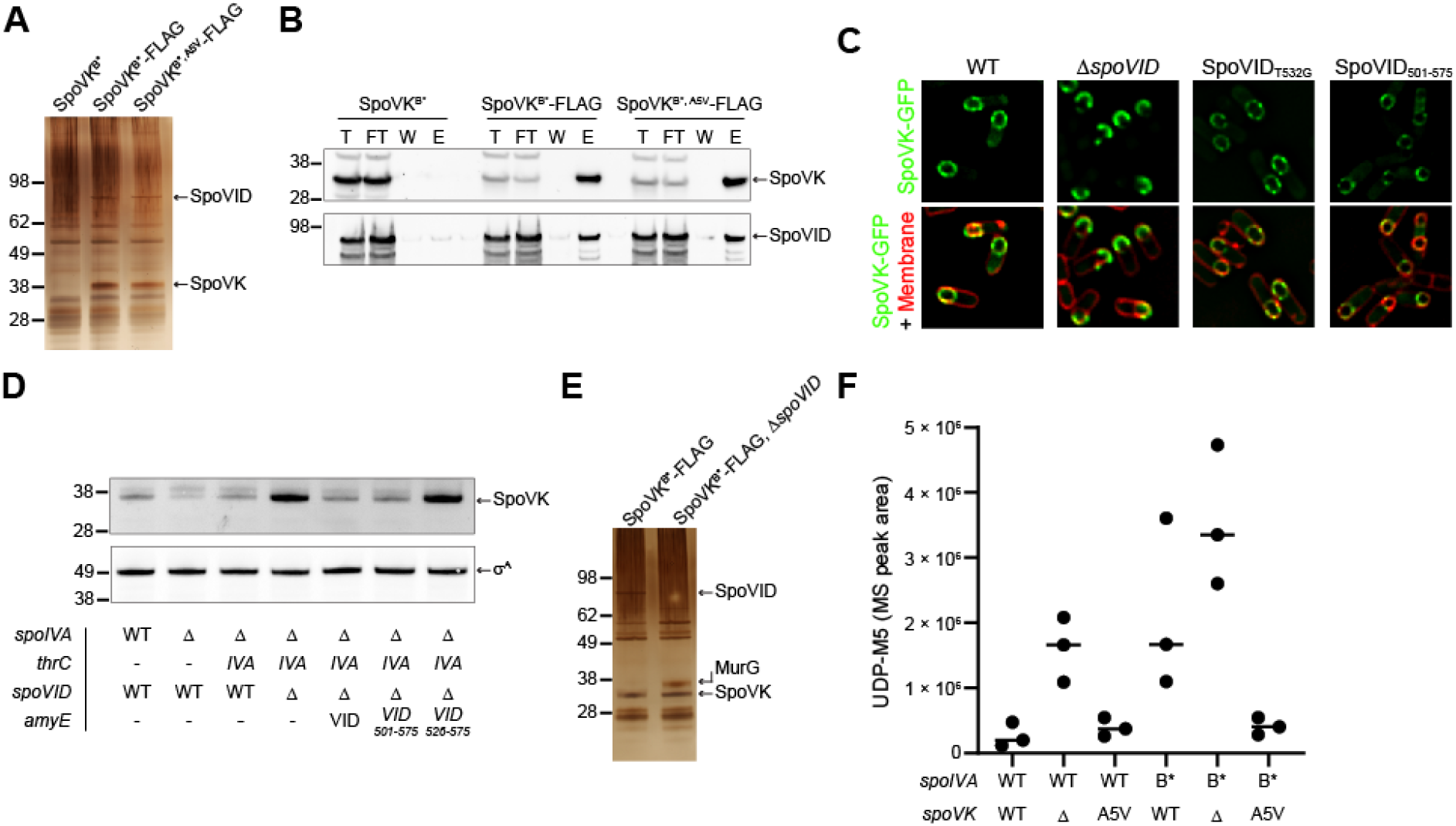
SpoVK interacts with SpoVID and MurG. (A) Silver stain SDS-PAGE of the elution fraction of the SpoVK^B^*-Flag immunoprecipitation from cells harvested 4 h after the induction of sporulation with anti-FLAG magnetic beads. Strains: TD883, TD884, and TD1193. (B) Immunoblot of Total (T), Flow Through (FT), Wash (W), and Elution (E) of the SpoVK^B^*-Flag immunoprecipitation from cells harvested 4h after the induction of sporulation with anti-Flag magnetic beads using anti-SpoVK and anti-SpoVID antisera. Strains: TD883, TD884, TD1193. (C) Subcellular localization of SpoVK-GFP in the presence or absence of SpoVID, or in cells producing SpoVID containing a defective LysM domain (T532G), or the SpoVID LysM domain alone (residues 501-575) 3.5 h after the induction of sporulation (Strains TD604, TD651, TD695, and TD1257). (D) Immunoblot of cell extracts of sporulating *B. subtilis* using anti-SpoVK or anti-σ^A^ antisera, from cells harvested at 4h. (Strains: PY79, KP73, KR394, JB171, JB174, TD892, and TD893). (E) Silver stain SDS-PAGE of the elution fraction of the SpoVK^B^*-Flag immunoprecipitation from Δ*spoVID* cells harvested 4 h after the induction of sporulation with anti-FLAG magnetic beads. Strains TD884 and TD1268. (F) Accumulation of peptidoglycan precursor Park’s nucleotide in strains of *B. subtilis* harboring WT (lanes 1-3) or defective (lanes 4-6) alleles of *spoIVA*, in the presence or absence (lanes 2, 5) of *spoVK* or the presence of *spoVK*^A5V^ (lanes 3, 6).

To identify other interacting partners for SpoVK, we repeated the immunoprecipitation of SpoVK^B*^-FLAG, but this time in the absence of SpoVID, which revealed a ∼38 kDa co-purifying band that was identified as MurG (Fig. 4E). MurG is the glycosyltransferase that catalyzes the formation of lipid II by transferring the UDP-activated GlcNac to the C4 hydroxyl of MurNAc in lipid I (37). Previously, we had reported that SpoVID sequesters lipid II in response to coat assembly defects. Hence, given the evolutionary affinities of SpoVK (Fig. 3B-C), we wondered if SpoVK was also involved in this pathway, perhaps as a chaperone that regulates MurG activity and thereby the amount of lipid II produced during sporulation. To test this, we examined how the presence or absence of SpoVK influences the accumulation of the peptidoglycan precursor Park’s nucleotide, which is the immediate precursor for lipid I synthesis (the substrate for MurG). In an otherwise WT cell 5.5 h after induction of sporulation, deletion of *spoVK* resulted in increased accumulation of Park’s nucleotide, indicating a likely defect in lipid I or lipid II synthesis (Fig. 4F, lanes 1-2). A similar accumulation in Park’s nucleotide was not observed in the presence of the *spoVK*^A5V^ (Fig. 4F, lane 3). When coat assembly is impaired by an ATPase-defective SpoIVA (SpoIVA^B*^) that does not polymerize, we observed an increase in the accumulation of Park’s nucleotide, similar to what we observed previously (28). This accumulation was exacerbated in the absence of SpoVK but was corrected in the presence of SpoVK^A5V^ (Fig. 4F, lanes 4-6). In sum, the data thus far are consistent with a model in which SpoVK positively influences MurG to promote cortex assembly. Further, the data suggest that SpoVID, via its C-terminal LysM domain, negatively regulates SpoVK activity, albeit independently of its lipid II-binding function.

### The forespore surface is more acidic relative to the mother cell cytoplasm

To test if the negative regulatory activity of the SpoVID LysM domain on SpoVK is via a direct interaction, we immunoprecipitated SpoVK^B*^-FLAG from cell extracts 4 h after induction of sporulation using cells that also produced GFP fused to the LysM domain of SpoVID (GFP-SpoVID^501-575^). Immunoblotting the fractions using anti-GFP antibodies indicated that GFP-SpoVID^501-575^ specifically co-purified with SpoVK^B*^-FLAG (Fig. 5A). Since the *spoVK*^A5V^ suppressor mutation mapped to the N-terminal domain, which is predicted to play a role in recruiting the substrate, we next sought to identify which protein binds to the SpoVK N-terminus. We therefore immunoprecipitated a SpoVK^B*^-FLAG variant in which amino acids 2-5 were deleted (SpoVK^Δ2-6^-FLAG) or harbored the A5V suppressor substitution (SpoVK^A5V^-FLAG) from cell extracts 4 h after induction of sporulation and first monitored the co-purification of SpoVID. Immunoprecipitation of SpoVK ^Δ2-6^-FLAG or SpoVK-FLAG resulted in similar co-purification of

**Figure 5.**
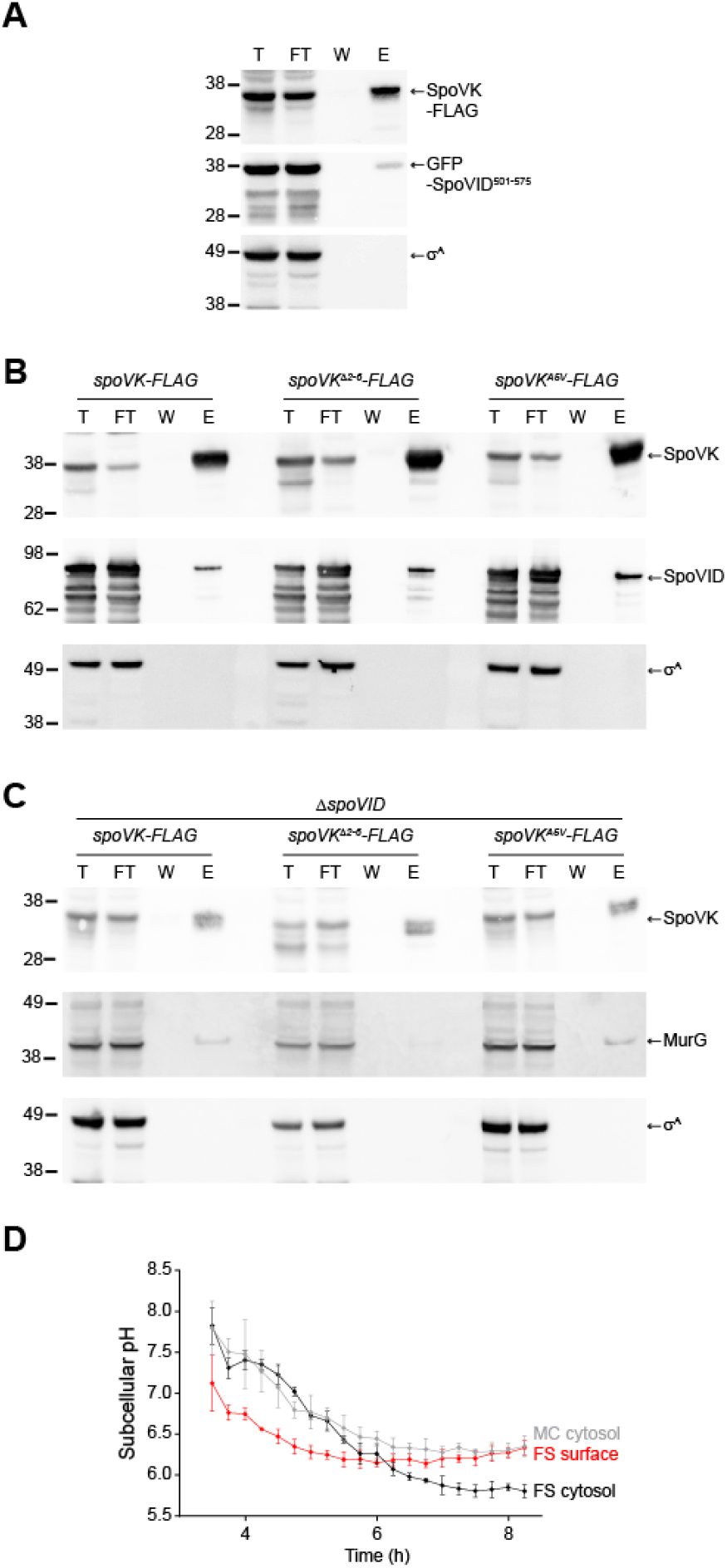
SpoVK interacts with SpoVID and MurG at an acidified forespore surface. (A) Immunoblots of total (T), flow through (FT), wash (W), and elution (E) fractions of the SpoVK^B^*-Flag immunoprecipitation from cells producing GFP-SpoVID^501-575^ (containing the SpoVID LysM domain) harvested 4h after the induction of sporulation with anti-Flag magnetic beads using anti-SpoVK, anti-GFP, or anti-σ^A^ antisera. Strain: TD1281. (B-C) Immunoblots of fractions described above, of the SpoVK^B^*-Flag immunoprecipitation (left), SpoVK^B^*-Flag with residues 2-5 deleted (middle), or SpoVK^B^*-Flag harboring the A5V substitution from cells that either (B) produce or (C) do not produce SpoVID, using anti-SpoVK, anti-SpoVID, or anti-σ^A^ antisera. Strains: TD1267, TD1277, TD1283, TD1268, TD1278, and TD1284. (D) Subcellular pH measurements in a sporangium near the producing IpHluorin fusions to SpoVM (to measure the forespore surface, “FS surface”, red), or free IpHluorin produced in the mother cell cytosol (“MC cytosol”, gray), or forespore cytosol (“FS cytosol”, black) taken at indicated time points after induction of sporulation. pH values were obtained by measurement of the emission fluorescence at 510 nm after excitation at 390 nm and 470 nm. After obtaining the emission ratio of 390 nm/470 nm, pH was calculated using a calibration curve obtained by growing IpHluorin-producing cells in media of defined pH in the presence of an electrochemical gradient dissipator. Data points represent mean of 4 independent biological replicates; errors are S.D. Strains: SC765, SC767, and SC766.

SpoVID (Fig. 5B), indicating that the N-terminus of SpoVK is not necessary for the interaction of SpoVK with SpoVID. We then repeated the experiment in a strain harboring a deletion of *spoVID*, which increased the interaction of SpoVK with MurG (Fig. 4E). An Alphafold2 multimer model suggested that the N-terminal domain of SpoVK is likely to form part of its interaction interface with MurG (Fig. S1E). Consistent with this, whereas MurG co-purified with SpoVK-FLAG and SpoVK^A5V^-FLAG, the amount of MurG that co-purified with SpoVK ^Δ2-6^-FLAG was diminished (Fig. 5C), indicating that the N-terminus of SpoVK is required for the interaction between SpoVK and MurG. In conclusion, although SpoVID interacts with SpoVK, SpoVID is likely not a substrate for the SpoVK. However, since the interaction of MurG with SpoVK depends on the N-terminus of SpoVK, we conclude that MurG is likely a substrate of the SpoVK chaperone.

MurG is an essential protein that reportedly does not require activation by a chaperone for normal function (37), so we wondered why it would need to be activated during sporulation. At the onset of sporulation, MurG redeploys from the mother cell membrane to the outer forespore membrane (30). Previously, based on two observations, we speculated that the forespore surface may represent a unique nanoenvironment in the mother cell cytosol. First, the LysM domain of SpoVID displays an unusually low pI (4.8) and is only functional in vitro in acidic buffer conditions (28). Second, SpoIVA, which is among the most abundant proteins on the forespore surface (14), also displays an acidic isoelectric point (4.5). Previous studies have shown that the forespore interior is ultimately acidified compared to the mother cell cytosol, which remains largely at neutral pH (38, 39), but the status of the nanoenvironment immediately atop the developing forespore in the mother cell cytosol is not known. To measure the pH of this region, we fused the pH-sensitive fluorescent protein IpHluorin to SpoVM, a peripheral membrane protein that tethers SpoIVA to the forespore surface (40). As controls, we also measured the pH of the mother cell and forespore cytosol by producing free IpHluorin either under control of a mother cell-specific promoter (*P*_spoVM_) or forespore-specific promoter (*P*_spoIIQ_). All Iphlorin fusions localized to the expected subcellular location (Fig. S3A-D). At the onset of our measurements, 3.5 h after induction of sporulation, the forespore and mother cell cytosols displayed a similar pH: 7.8 ± 0.33 and 7.8 ± .23, respectively, but the nanoenvironment immediately surrounding the forespore surface displayed a pH of 7.1 ± 0.34. At 4.25 h after induction of sporulation, the overall pH of both the mother cell (7.3 ± 0.3) and forespore (7.4 ± 0.07) cytosols gradually decreased, but remained similar to one another; however, the forespore surface pH was further reduced (6.6 ± 0.02). The relatively lower pH of the forespore surface persisted until 6 h after induction of sporulation, when the forespore cytosol became acidified to a similar extent as the forespore surface. At 8 h after induction of sporulation, the mother cell cytosol (6.35 ± 0.13) was similar to that of the forespore surface (6.33 ± 0.08), whereas the cytosol of the forespore was further acidified (5.8 ± 0.08), as previously reported (38, 39). We therefore conclude that the mother cell cytosol is not uniform and that, even in the absence of membrane-bound compartments, certain patches of the cytosol can exhibit unique chemical and physical properties.

## DISCUSSION

In non-spherical bacteria such as rods, peptidoglycan assembly at differently shaped regions of the cell (for example, the lateral edge versus the division septum) relies on a shared pool of peptidoglycan precursors in the cytosol that are differently assembled, depending on where on the cell surface the precursors need to be incorporated. The differential utilization of this precursor pool is achieved by specialized complexes (the “divisome” at septa or the “elongasome” for cylindrical growth) that permit the coordination of cell wall synthesis with cell growth.

The sporulation program typically employs factors that are exclusively used for this pathway, even if that requires duplicating an existing gene and modifying it for sporulation (41, 42). Given the precedence of shared peptidoglycan precursor usage for the assembly of different cell wall material, it is interesting that a sporulation-specific peptidoglycan assembly complex has evolved to generate the specialized cortex, which nonetheless utilizes the same pool of peptidoglycan precursors as used for vegetative growth. Indeed, during sporulation, the widely conserved MurG protein, which synthesizes the lipid II precursor, redeploys from the mother cell plasma membrane, where it participates in constructing the vegetative cell wall that surrounds the mother cell, onto the outer forespore membrane during sporulation, where it participates in assembling the cortex surrounding the developing forespore (30). In this study, we propose that the redeployed MurG additionally requires a sporulation-specific chaperone, the previously uncharacterized SpoVK protein, to perform its glycosyltransferase function at the forespore surface. We suggest that this requirement arises from a nanoenvironment immediately surrounding the forespore, which we determined was more acidic than the rest of the mother cell cytosol. The requirement of SpoVK during sporulation also conveniently provides an additional step of regulating cortex assembly. Previously, we demonstrated that cortex assembly is linked to proper initiation of coat assembly by a checkpoint mechanism that actively monitors the polymerization state of the spore coat basement layer: when the coat mis-assembles, cortex assembly is inhibited by sequestration of lipid II by SpoVID (43). However, it was unclear if additional lipid II synthesis would be downregulated in this circumstance to prevent overwhelming this lipid II sequestration mechanism. In our updated model (Fig. 6), we propose that upon sensing a coat assembly defect, SpoVID also inhibits SpoVK by directly binding SpoVK via the LysM domain of SpoVID. Thus, SpoVID prevents the activation of MurG at the forespore surface, resulting in reduced flux of lipid II through the cortex biosynthesis pathway.

**Figure 6.**
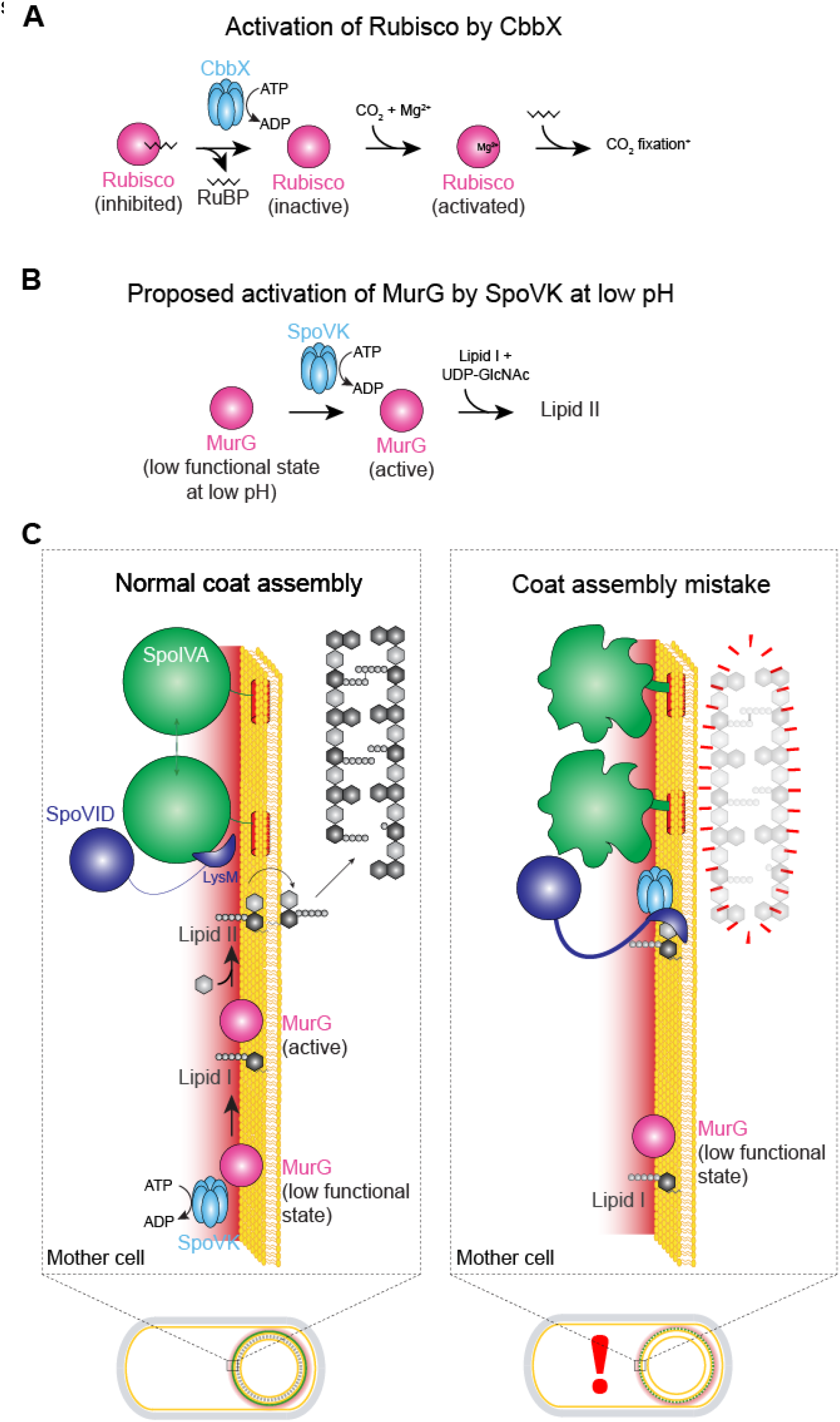
Model for the regulated activation of MurG by SpoVK. (A) Activation of Rubisco, which catalyzes fixation of CO2 in photosynthesis, by AAA+ chaperone CbbX. Rubisco (pink) forms an inactive complex with its substrate ribulose 1,5-bisphosphate (RuBP, black), which can be reactivated by CbbX (blue). Rubisco is subsequently activated by reacting with CO2 and Mg^2+^ (adapted from Mueller-Cajar *et al*. (1)). (B) Proposed activation of MurG by AAA+ chaperone SpoVK. MurG exists in a low functional state at the forespore surface due to the low pH in that nanoenvironment. SpoVK, which localizes to the forespore surface, helps fold MurG properly so that it may catalyze the conversion of lipid I to lipid II. (C) Depicted are a sporulating cell of *B. subtilis* that is wild type (left) or a mutant (right) that mis-assembles the spore coat. Expansions of the developing spore envelope (cortex, outer forespore membrane, and coat basement layer) are depicted above each cell. Low pH nanoenvironment is depicted as a red gradient. When SpoIVA (green) polymerizes properly, the LysM domain of SpoVID (purple) is occluded, permitting 1) lipid II to flip to the intermembrane space to incorporate into the assembling cortex and 2) SpoVK (blue) to activate MurG, thereby ensuring a steady flux of lipid II. When SpoIVA mis-assembles (right), the LysM domain of SpoVID is liberated, resulting in 1) sequestration of lipid II and 2) inhibition of SpoVK, resulting in reduced buildup of lipid II in the mother cell cytosol, due to the low functional state of MurG in the forespore surface nanoenvironment.

Our model for how SpoVID controls the coat assembly checkpoint by monitoring SpoIVA polymerization, sequestering lipid II, and modulating the activity of SpoVK is consistent with multiple in vivo and in vitro observations. First, SpoVK co-localizes on the forespore surface with MurG; in the absence of SpoVK, cortex peptidoglycan assembly is diminished and the resulting cell wall surrounding the forespore resembles that resulting from the deletion of other cortex morphogenesis factors (34, 44, 45). Notably, though, while other well-characterized cortex morphogenic proteins localize in the intermembrane space of the forespore, SpoVK is in the mother cell, where it can interact with peptidoglycan precursors, before lipid II is flipped across the outer forespore membrane. Interestingly, the undetectability of SpoVK at an intermediate time point during sporulation that coincides with the completion of cortex assembly and the dehydration of the spore core also suggests that SpoVK plays a defined role in cortex assembly. Second, the evolutionary relationship of SpoVK to the CbbX chaperone, which activates RuBisCo (1), suggests that SpoVK could also act as a comparable chaperone. RuBisCo is a large multimeric complex that is inhibited by its own substrate ribulose 1,5-bisphosphate (RuBP; Fig. 6A). CbbX, in contrast, is stimulated by RuBP, which CbbX binds via the C-terminal four-helical bundle characteristic of AAA+ ATPases. Thus, CbbX captures the C-terminal tail of the RuBisCO large subunit and causes it to release the inhibitor RuBP. Interestingly, MurG has been recently described to also form a large hexameric complex (46). Hence, like RuBisCO, a functional MurG complex might require assistance from a chaperone for assembly/activation in mutation in the putative substrate binding domain of SpoVK and observed that SpoVK interacts with MurG *in vitro* via this domain. Further, SpoVK shares sequence conservation with CbbX in the C-terminal four-helical bundle which binds the activating carbohydrate RuBP in the latter (Fig. S1B-C). Hence, it would also be of interest to explore if SpoVK might similarly be regulated by soluble carbohydrates such as substrates of MurG. Finally, we found that SpoVK is antagonized by SpoVID, likely by destabilization and that the C-terminal LysM domain of SpoVID, which stably interacts with SpoVK, is sufficient for this antagonism. Thus, detecting improper assembly of the spore coat, sequestering the lipid II peptidoglycan precursor, and reducing the further synthesis of lipid II is mediated by SpoVID, the central factor in the spore coat assembly checkpoint, that interacts with SpoIVA, SpoVK, and lipid II via a single domain.

Our findings have implications for the specific clade of AAA+ domains to which SpoVK belongs that was previously both poorly defined and understudied. The sporulation-specific interaction of SpoVK and MurG adds to the previous evidence from CbbX and EccA that members of this clade are dedicated chaperones that act narrowly on specific targets and/or in specific sub-cellular locations. Indeed, while most well-characterized members of the Classical AAA+ assemblage show fusions or direct physical interaction with diverse peptidase domains, we found no evidence for such associations in this clade. Instead, they typically show a diversity of N-terminal domains which might help recruit distinct substrates to an otherwise well-conserved AAA+ domain. Thus, rather than aiding protein degradation through unfolding, these chaperones might aid macromolecular assembly and activation in several as yet unstudied contexts. Interestingly, in addition to its role in secretion via T7SS, the mycobacterial EccA protein EccA1 has also been shown to be necessary for production of mycolic acids, key component of the actinobacterial cell envelope lipid. This role of EccA1 has been proposed to operate via potential chaperone action on several mycolic acid biosynthesis enzymes (47). Thus, multiple members of this clade might regulate cell envelope composition by acting on the cytosolic enzymes needed for their synthesis.

Despite the diminutive size of the cytoplasm of most bacteria, the subcellular organization of macromolecules in this compartment has become increasingly evident in the last several decades (48). Interestingly, our studies in understanding a bacterial developmental checkpoint led us to appreciate that cytosolic compartmentalization may also exist at a chemical level in the form of varying pH, which can necessitate specialized chaperones at distinct regions to carry out cellular functions. More generally, this raises the interesting possibility that the cytoplasm may not be uniform with respect to other chemical properties, such as ionic strength or oxidation state, as well. Perhaps discovering the subcellular localization of chaperones in other systems may highlight other sub-cytosolic locations that harbor chemically distinct regions.

## ACKNOWLEDGEMENTS

We thank members of the KSR lab for suggestions and comments on the manuscript; S. Gottesman, S. Wickner, A. Khare, T. Bauer, and M. Maurizi for discussions; J. Barriga for strains; F. Soheilian and C. Burks (CCR) for TEM sample preparation and imaging; and the CCR Genomics Core Facility for whole genome sequencing. This work was funded by the National Institutes of Health (NIH) grant R01GM138630 (D.L.P.), the Swedish Research Council, the Laboratory of Molecular Infection Medicine Sweden (MIMS), Umeå University, the Knut and Alice Wallenberg Foundation (KAW), the Kempe Foundation (F.C.), the Intramural Research Program of the NIH, the National Cancer Institute, the Center for Cancer Research (K.S.R.), and the National Library of Medicine (L.A.). This work utilized the NIH HPC Biowulf computer cluster (V.A. and L.A.).

## EXPERIMENTAL PROCEDURES

### Strain construction

Strains are otherwise isogenic derivatives of *B. subtilis* PY79 (49). Genes of interest were PCR amplified to include their native promoter and cloned into integration vectors pDG1662 (for insertion into the *amyE* locus), pDG1731(*thrC* locus), pSac-Cm (*sacA* locus), or pPyr-Cm (*pyrD* locus) (50, 51). Site-directed mutagenesis was performed using the QuikChange kit (Agilent). For pH measurements, 500 bp upstream of *spoIIQ* or *spoVM* genes were cloned upstream of *Iphluorin* and cloned into pDG1662 (51) to drive production of free Iphluorin in the forespore or mother cell cytosol, respectively. Alternatively, the spoVM ORF, downstream of its native promoter, was fused to *Iphluorin* (52) and cloned into pDG1662 to drive production of membrane-bound SpoVM-Iphluorin. Finally, IPTG-inducible *Iphluorin* was constructed by cloning Iphluorin into vector pDP150 (53). Resulting plasmids were integrated into the specified ectopic locus in the *B. subtilis* chromosome by double recombination.

### General methods

Sporulation efficiencies were calculated by growing cells in Difco Sporulation Medium (KD Medical) for at least 24 h. Nonsporulating cells and defective spores were killed by exposure to 80 °C for 20 min. Heat-killed cultures were serially diluted and colony forming units (cfu) of surviving cells were enumerated and reported relative to cfu enumerated in a parallel culture of WT (PY79) strain. To obtain spontaneous suppressor mutants, strain JPC221 (which harbors the *spoIVA*^T^* allele as the only copy of *spoIVA* and is unable to sporulate efficiently) was subjected to multiple rounds of growth and heat treatment in DSM to enrich for colonies that displayed increased heat resistance. The suppressor mutation was identified using whole genome sequencing as described previously (28). SpoVK, σ^A^, SpoVID, GFP, and MurG levels were by immunoblotting cell extracts prepared as described previously (36) using rabbit antiserum raised against recombinant, purified SpoVK-His_6_, σ^A^-His_6_, SpoVID-His_6_, GFP-His_6_, and MurG-His_6_ (Covance) as primary antibody and goat Starbright Blue 700 (Bio-Rad) as secondary antibody.

### Sequence Analysis

Sequence similarity searches were performed using the PSI-BLAST program with a profile-inclusion threshold set at an e-value of 0.01 (54). The searches were conducted against the NCBI non-redundant (nr) database, or the same database clustered down to 50% sequence identity using the MMseqs program, or a curated database of 7423 representative genomes from across the tree of life. Profile-profile searches were performed with the HHpred program (55, 56). Multiple sequence alignments (MSAs) were constructed using the FAMSA and MAFFT programs (57, 58). Sequence logos were constructed using these alignments with the ggseqlogo library for the R language (59).

### Structure Analysis

PDB coordinates of structures were retrieved from the Protein Data Bank and structures were rendered, compared, and superimposed using the Mol* program (60). Structural models were generated using the AlphaFold2 and AlphaFold-Multimer programs (61, 62). Multiple alignments of related sequences (>30% similarity) were used to initiate HHpred searches for the step of identifying templates to be used by the neural networks deployed by these programs.

### Comparative Genomics and Phylogenetic Analysis

Clustering of protein sequences and the subsequent assignment of sequences to distinct families was performed by the MMseqs program (63), adjusting the length of aligned regions and bit-score density threshold empirically. Phylogenetic analysis was performed using the maximum-likelihood method with the IQTree program (64) and multiple protein substitution models such as Dayhoff, Poisson, and JTTMutDC. The FigTree program (http://tree.bio.ed.ac.uk/software/figtree/) was used to render phylogenetic trees. Gene neighborhoods were extracted through custom PERL scripts from genomes retrieved from the NCBI Genome database.

### Epifluorescence microscopy

*B. subtilis* cells were induced to sporulate by the resuspension method in SM medium (65). At indicated time points, 100 µl of culture was harvested and resuspended in 10 µl SM containing 5 µg ml^-1^ FM4-64 fluorescent dye to visualize membranes. Resuspensions were then placed on a glass-bottom culture dish (MatTek) and covered with a 1% agarose pad made with SM. Cells were viewed at room temperature with a DeltaVision Core microscope system (Applied Precision/GE Healthcare). Seven planes were acquired every 200 nm and the data were deconvolved using SoftWorx software as described previously (66). Additional image adjustments were performed using Fiji software.

### Transmission electron microscopy

*B. subtilis* cells were induced to sporulate by resuspension in SM for the indicated time period (> 24 h for mature spores), harvested by centrifugation, washed with water, and fixed in 4% formaldehyde and 2% glutaraldehyde in 0.1 M cacodylate buffer, post fixed using a 1% osmium tetroxide solution, then dehydrated sequentially in 35%, 50%, 75%, 95% and 100% ethanol followed by 100% propylene oxide. Cells were infiltrated in an equal volume of 100% propylene oxide and epoxy resin overnight and embedded in pure resin the following day. The epoxy resin was cured at 55 °C for 48 h. The cured block was thin-sectioned and stained in uranyl acetate and lead citrate. The sample was imaged with a Hitachi H7600 TEM equipped with a CCD camera (67).

### SpoVK purification

SpoVK-His_6_ was purified by affinity chromatography under denaturing conditions and renatured. Overnight cultures of *E. coli* strains BL21(DE3) pTD211 (expressing *spoVK-His_6_*) or pTD308 (expressing *spoVK^D162A,E163A^-His_6_*) in LB containing 50 µg ml^-1^ kanamycin for plasmid maintenance were diluted 1:100 into 500 ml LB/kanamycin and grown at 37 °C shaking at 250 rpm for 2 h. Isopropyl thiogalactopyranoside was then added at 1 mM final concentration to induce protein production and cells were grown for a further 3 h. Cells were harvested by centrifugation and cell pellets were stored at -80 °C. Cell pellets were resuspended in lysis buffer (100 mM sodium phosphate, 10 mM Tris-HCl, 8 M Urea at a final pH of 7.5) and disrupted in a French pressure cell at ca. 20,000 psi. Insoluble material was removed by centrifugation at ∼100,000 × g, and the supernatant was applied onto a 1.5 ml (bed volume) Ni^2+^-NTA agarose column equilibrated with lysis buffer. The column was washed with 30 ml of lysis buffer and eluted with 4 ml lysis buffer containing 250 mM imidazole. 3 ml of purified SpoVK-His_6_ was then dialyzed once overnight against 500 ml renaturation buffer (500 mM L-arginine, 100 mM NaCl at pH 7.5) using a 10 kDa MWCO membrane. Dialyzed sample was then collected, and insoluble material was removed by centrifugation at ∼15,000 × g for 5 min at 4 °C. 500 µl of the supernatant was then applied to a Superdex 200 Increase size exclusion chromatography column (Cytiva) and separated using renaturation buffer using flow rate of 0.75 ml min^-1^. The final peak, corresponding to an approximate trimer of SpoVK, was used for analysis.

### Co-immunoprecipitation

*B. subtilis* cells were induced to sporulate by resuspension in 15 ml SM for 4 h, and cell pellets harvested by centrifugation were stored at -80 °C. Pellets were resuspended in 1 ml protoplast buffer (0.5 M sucrose, 20 mM MgCl_2_, 10 mM potassium phosphate at pH 6.8, 1 mg/ml lysozyme) and incubated at 37 °C for 25 min to generate protoplasts. Protoplasts were harvested by centrifugation and resuspended in 1 ml binding buffer (50 mM Tris-HCl at pH 7.5, 150 mM NaCl, 7.5% glycerol, 0.2% Triton-X-100). Cell debris was removed by centrifugation at ∼15,000 × g, and the supernatant was combined with an additional 150 µl of binding buffer and added to 50 µl magnetic agarose beads (Pierce) equilibrated with binding buffer and incubated at 4 °C for 1 h. Beads were washed thrice with 1 ml of binding buffer and eluted with 100 mM glycine at pH 2.7.

### Preparation and analysis of forespore peptidoglycan

Cortex peptidoglycan was prepared and analyzed as previously described (68). Briefly, 50 ml cultures were induced to sporulate by resuspension as described above. 5 h after induction of sporulation, cells were harvested by centrifugation and pelleted cells were resuspended in 5 ml SMM protoplast solution (0.5 M sucrose, 20 mM Maleic Acid, 20 mM MgCl_2_) containing 25 mg/ml lysozyme to digest mother cell peptidoglycan and incubated at 37 °C for 15 min. 45 ml boiling lysis buffer (4% sodium dodecyl sulfate, 50 mM dithiothreitol) was then added to the resulting protoplasts and boiled for 20 min. After cooling to room temperature, insoluble material was collected by centrifugation at 21,000 × g for 30 min and resuspended in 1 ml water, boiled for 5 min to solubilize residual SDS, then centrifuged at 21,000 × g for 20 min to collect insoluble material. Washes were repeated until no SDS was detected. Resuspended material in 1 ml buffer (100 mM Tris at pH 7.0, 20 mM MgSO_4_) were treated with 10 µg/ml DNase I and 50 µg/ml RNase A for 2h at 37 °C. 100 µg trypsin and 10 mM CaCl_2_ were added and the sample was incubated at 37 °C for 16 h. Insoluble material was collected by centrifugation at 21,000 × g for 20 min, resuspended in 1 ml 1% SDS, and boiled for 20 min to inactivate trypsin. Samples were washed with water as described above and isolated spore peptidoglycan was digested with 125 U of Mutanolysin in a total volume of 250 µl of 12.5 mM NaPO_4_ at pH 5.5 for 16 h at 37 °C. Solubilized muropeptides were separated by reverse phase HPLC as described previously (69).

### Peptidoglycan accumulation

Quantification of Park’s nucleotide was performed as described previously (28). Briefly, cells were induced to sporulate by resuspension in SM, harvested at t = 5.5 h, and washed thrice with ice cold 0.9% NaCl, resuspended in a final volume of 100 µl 0.9% NaCl, and boiled for 5 min to extract soluble peptidoglycan precursors. Insoluble material was pelleted by centrifugation at 21,000 × g for 5 min and the resulting supernatant was filtered through a 0.22 µm pore-size filter. Detection and quantification of Park’s nucleotide was performed by LC-MS analysis using a UPLC system (Waters) equipped with an ACQUITY UPLC BEH C18 column (130Å pore size, 1.7 μm particle size, 2.1 mm x 150 mm, Waters) coupled to a Xevo G2-XS QTOF mass spectrometer (Waters). Chromatographic separation of the soluble fraction was achieved using a linear gradient from 0.1% formic acid in water to 0.1% formic acid in acetonitrile over 18 min at 45 °C. The QTOF instrument was operated in positive ionisation mode and detection of UDP-M5 was performed in the untargeted MS^e^ mode. The MS parameters were set as follows: capillary voltage 3 kV, source temperature 120 °C, desolvation temperature 350 °C, sample cone voltage 40 V, cone gas flow 100 L h^-1^, and desolvation gas flow 500 L h^-1^. Data acquisition and processing was performed using the UNIFI software (Waters). To quantify the Park’s nucleotide, its calculated [M+2H]^2+^ ion of *m/z* 597.68 was extracted from the total ion chromatogram, and the corresponding peak in the resulting extracted ion chromatogram was integrated to give a peak area.

### ATP hydrolysis assay

ATP hydrolysis was measured as previously described (16). Briefly, varying concentrations of ATP were incubated with 1 µM purified SpoVK-His_6_ or SpoVK^B*^-His_6_ in 100 µl of buffer (500 mM L-arginine at pH 7.5, 100 mM NaCl, 5 mM MgCl_2_) for 20 min at 37 °C. Concentration of inorganic phosphate was determined using the Malachite Green Phosphate Assay kit (BioAssay Systems) according to manufacturer’s protocol. Absorbance at 620 nm (Spark 10M plate reader, Tecan) of each reaction was compared to absorbances of known concentrations of phosphate standards. Absorbances from control reactions performed in the absence of SpoVK for each ATP concentration were subtracted from absorbances of the respective reactions with SpoVK to eliminate background hydrolysis. For each ATP concentration, hydrolysis rates were plotted using GraphPad Prism 7; *V*_max_ and *K*_m_ values were determined by fitting the data to Michaelis–Menten equation using best-fit values.

### Intracellular pH measurements

Strains SC765, SC766, and SC767 were induced to sporulate by the resuspension method as described above. 200 µl of each culture were then placed into individual wells of black-walled 96-well plates and placed in a Synergy H1 plate reader (BioTek) and grown at 37 °C, with continuous shaking. Fluorescence signal (emission at 510 nm, with excitation at 390 nm and 470 nm) was measured every 15 min and the 390/470 ratios were calculated. To convert the 390/470 ratios to pH values, we constructed a calibration curve as described previously (39) (Fig. S3E). Briefly, strain SC777, which produces IPTG-inducible Iphluorin, was grown in casein-hydrolysate (CH) medium with 1 mM IPTG, then induced to sporulate by resuspension in Sterlini-Mandelstam medium (65) containing 10 mM Tris at pH 7.5, 1 µM nigericin, and 1 µM valinomycin (to equilibrate external and internal pH) that was adjusted to different pH values ranging from 5 to 8.5 using 5N NaOH or 6N HCl. Fluorescence signals of cells grown at different pH values were measured as described above, and 390/470 ratios were calculated and plotted as a function of pH. The resulting plot was fit with a linear curve with equation y = 0.7868x – 2.638, where y represents 390/470 ratio and x represents pH.

## SUPPORTING INFORMATION

**Table S1.** Cortex peptidoglycan analysis of WT and Δ*spoVK* strains. Extracted peptidoglycan from developing forespores of the wild type and Δ*spoVK* strains harvested 5 h after the onset of sporulation was digested with mutanolysin and separated using HPLC. Muropeptides were identified based on previous studies of elution times^1,2^. Peaks were then integrated, and peptidoglycan structural parameters were calculated. Errors are S.D.

| PG species | WT | $\Delta spoVK$ |
| --- | --- | --- |
| % muramic acid as lactam | 33.3 ± 1.5 | 6.5 ± 3.0 |
| % muramic acid with alanine | 20.6 ± 0.6 | 19.5 ± 0.9 |
| % muramic acid with tripeptide | 15.9 ± 1.9 | 42.9 ± 3.7 |
| % muramic acid with tetrapeptide | 30.2 ± 0.4 | 31.2 ± 1.4 |
| % muramic acid with pentapeptide | 0.0 ± 0.0 | 0.0 ± 0.0 |
| % peptide as tetrapeptide | 65.6 ± 3.0 | 42.2 ± 2.5 |
| % peptide as tripeptide | 34.4 ± 3.0 | 57.8 ± 2.5 |
| % peptide in crosslinks | 19.0 ± 0.6 | 20.4 ± 2.2 |
| % peptide in effective crosslinks | 19.0 ± 0.6 | 20.4 ± 2.2 |
| % disaccharide units in disaccharides | 36.4 ± 3.0 | 87.1 ± 6.0 |
| % disaccharide units in tetrasaccharides | 54.7 ± 2.8 | 12.9 ± 6.0 |
| % disaccharide units in hexasaccharides | 9.0 ± 0.5 | 0.0 ± 0.0 |
| % lactam in regular distribution | 82.1 ± 0.9 | 100.0 ± 0.0 |
| % disaccharide peptide crosslinked | 23.8 ± 0.3 | 20.4 ± 2.2 |
| % tetrasaccharide peptide crosslinked | 5.0 ± 0.2 | 0.0 ± 0.0 |
| % lactam reduced | 0.2 ± 0.0 | 0.2 ± 0.0 |
| % tripeptide crosslinked | 39.6 ± 0.8 | 31.1 ± 1.7 |
| % tetrapeptide crosslinked | 8.2 ± 0.5 | 5.7 ± 4.7 |
| % disaccharide per effective crosslink | 11.5 ± 0.7 | 6.6 ± 0.4 |
| % muramic acid with crosslink | 8.8 ± 0.5 | 15.1 ± 0.9 |

<sup>1</sup>DOI: 10.1128/JB.182.16.4491-4499.2000
<sup>2</sup>DOI: 10.1128/jb.178.22.6451-6458.1996

**Table S2.**
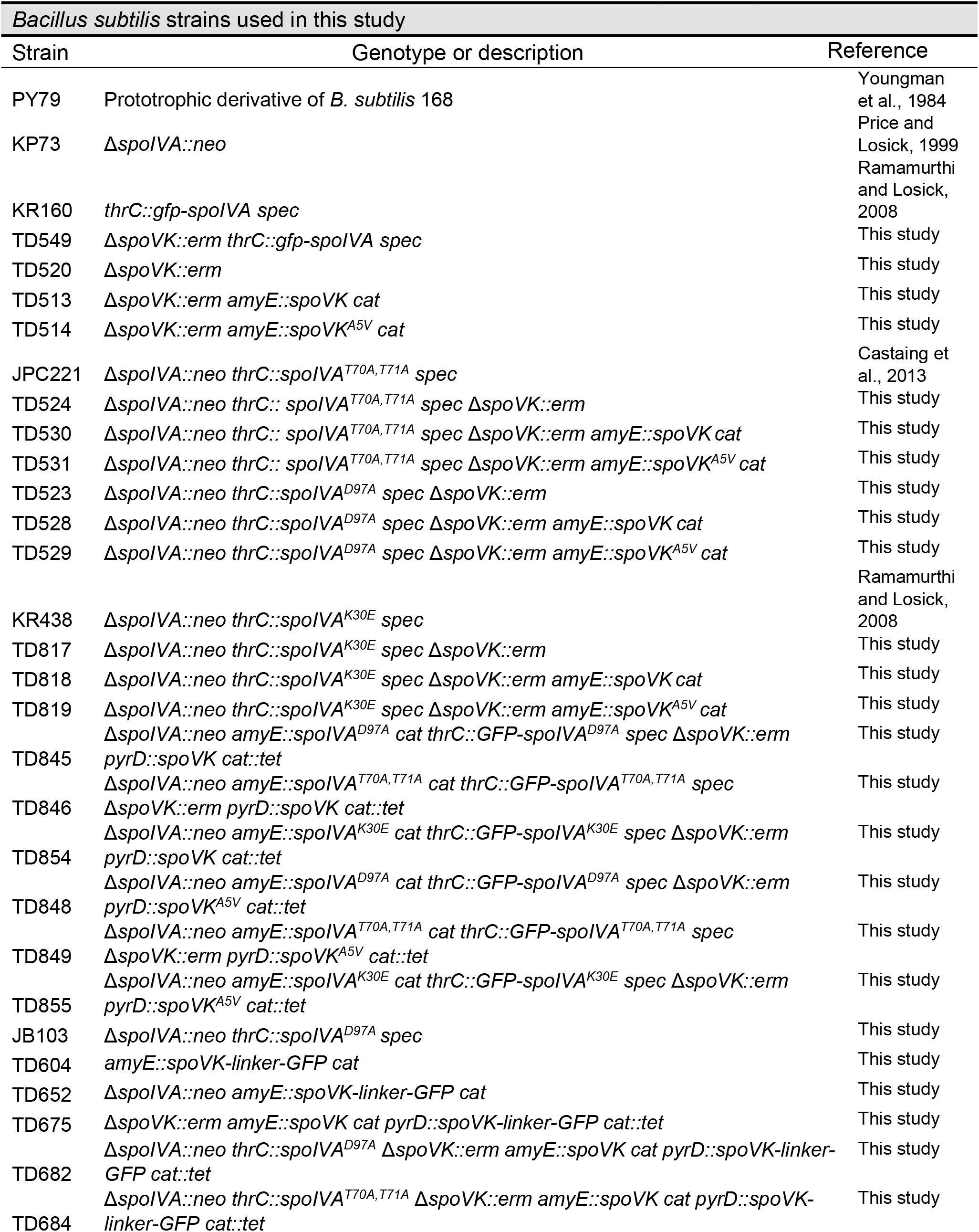
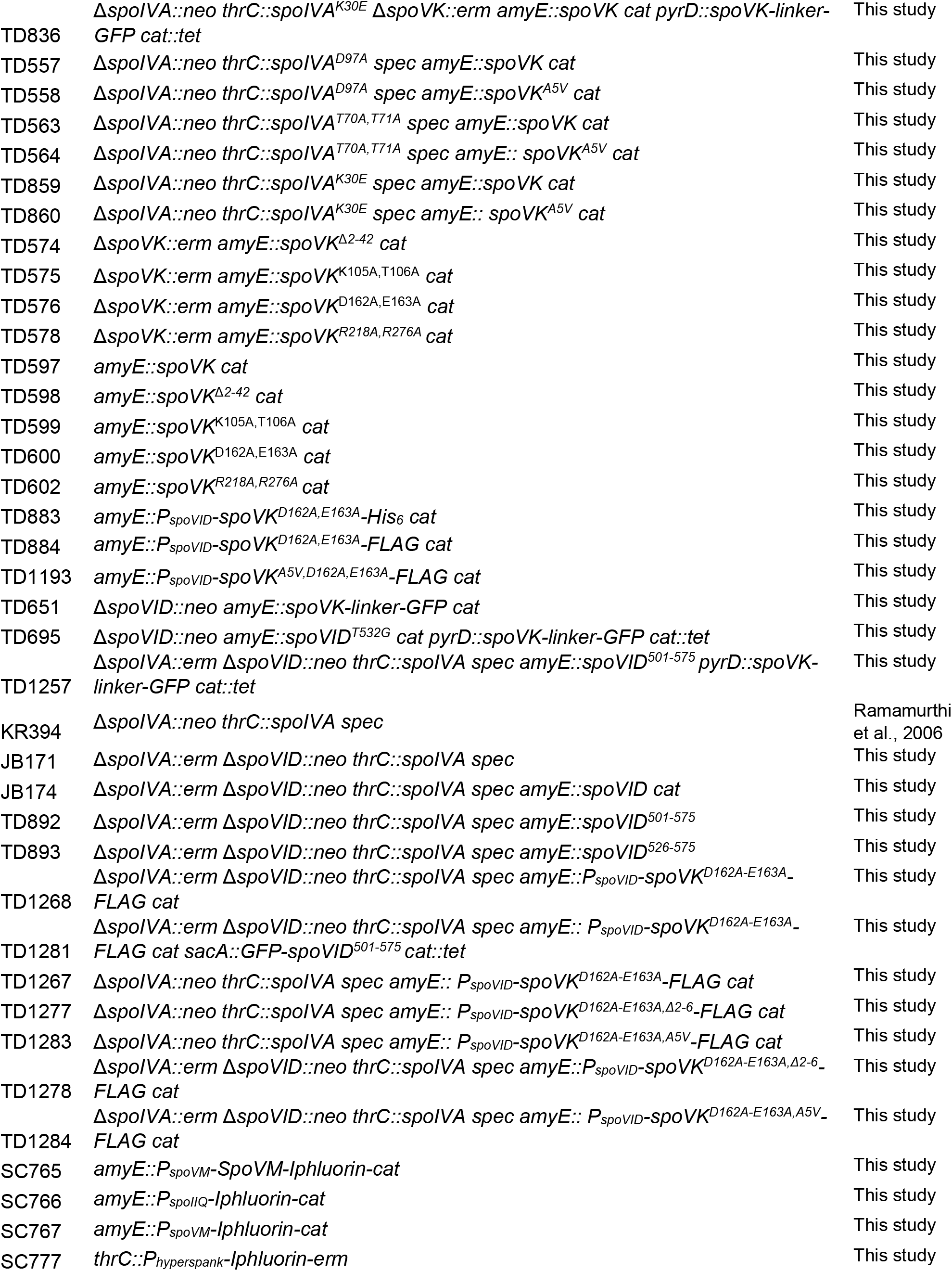
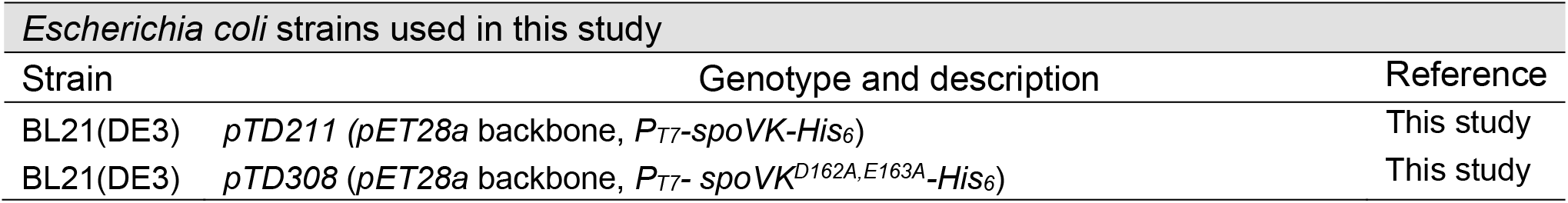
Bacterial Strains.

| *Bacillus subtilis* strains used in this study |  |  |
| --- | --- | --- |
| Strain | Genotype or description | Reference |
| PY79 | Prototrophic derivative of *B. subtilis* 168 | Youngman et al., 1984 |
| KP73 | $\Delta spoIVA::neo$ | Price and Losick, 1999 |
| KR160 | *thrC::gfp-spoIVA spec* | Ramamurthi and Losick, 2008 |
| TD549 | $\Delta spoVK::erm thrC::gfp-spoIVA spec$ | This study |
| TD520 | $\Delta spoVK::erm$ | This study |
| TD513 | $\Delta spoVK::erm amyE::spoVK cat$ | This study |
| TD514 | $\Delta spoVK::erm amyE::spoVK^{A5V} cat$ | This study |
| JPC221 | $\Delta spoIVA::neo thrC::spoIVA^{T70A,T71A} spec$ | Castaing et al., 2013 |
| TD524 | $\Delta spoIVA::neo thrC::spoIVA^{T70A,T71A} spec \Delta spoVK::erm$ | This study |
| TD530 | $\Delta spoIVA::neo thrC::spoIVA^{T70A,T71A} spec \Delta spoVK::erm amyE::spoVK cat$ | This study |
| TD531 | $\Delta spoIVA::neo thrC::spoIVA^{T70A,T71A} spec \Delta spoVK::erm amyE::spoVK^{A5V} cat$ | This study |
| TD523 | $\Delta spoIVA::neo thrC::spoIVA^{D97A} spec \Delta spoVK::erm$ | This study |
| TD528 | $\Delta spoIVA::neo thrC::spoIVA^{D97A} spec \Delta spoVK::erm amyE::spoVK cat$ | This study |
| TD529 | $\Delta spoIVA::neo thrC::spoIVA^{D97A} spec \Delta spoVK::erm amyE::spoVK^{A5V} cat$ | This study |
| KR438 | $\Delta spoIVA::neo thrC::spoIVA^{K30E} spec$ | Ramamurthi and Losick, 2008 |
| TD817 | $\Delta spoIVA::neo thrC::spoIVA^{K30E} spec \Delta spoVK::erm$ | This study |
| TD818 | $\Delta spoIVA::neo thrC::spoIVA^{K30E} spec \Delta spoVK::erm amyE::spoVK cat$ | This study |
| TD819 | $\Delta spoIVA::neo thrC::spoIVA^{K30E} spec \Delta spoVK::erm amyE::spoVK^{A5V} cat$ | This study |
| TD845 | $\Delta spoIVA::neo amyE::spoIVA^{D97A} cat thrC::GFP-spoIVA^{D97A} spec \Delta spoVK::erm pyrD::spoVK cat::tet$ | This study |
| TD846 | $\Delta spoIVA::neo amyE::spoIVA^{T70A,T71A} cat thrC::GFP-spoIVA^{T70A,T71A} spec \Delta spoVK::erm pyrD::spoVK cat::tet$ | This study |
| TD854 | $\Delta spoIVA::neo amyE::spoIVA^{K30E} cat thrC::GFP-spoIVA^{K30E} spec \Delta spoVK::erm pyrD::spoVK cat::tet$ | This study |
| TD848 | $\Delta spoIVA::neo amyE::spoIVA^{D97A} cat thrC::GFP-spoIVA^{D97A} spec \Delta spoVK::erm pyrD::spoVK^{A5V} cat::tet$ | This study |
| TD849 | $\Delta spoIVA::neo amyE::spoIVA^{T70A,T71A} cat thrC::GFP-spoIVA^{T70A,T71A} spec \Delta spoVK::erm pyrD::spoVK^{A5V} cat::tet$ | This study |
| TD855 | $\Delta spoIVA::neo amyE::spoIVA^{K30E} cat thrC::GFP-spoIVA^{K30E} spec \Delta spoVK::erm pyrD::spoVK^{A5V} cat::tet$ | This study |
| JB103 | $\Delta spoIVA::neo thrC::spoIVA^{D97A} spec$ | This study |
| TD604 | $amyE::spoVK-linker-GFP cat$ | This study |
| TD652 | $\Delta spoIVA::neo amyE::spoVK-linker-GFP cat$ | This study |
| TD675 | $\Delta spoVK::erm amyE::spoVK cat pyrD::spoVK-linker-GFP cat::tet$ | This study |
| TD682 | $\Delta spoIVA::neo thrC::spoIVA^{D97A} \Delta spoVK::erm amyE::spoVK cat pyrD::spoVK-linker-GFP cat::tet$ | This study |
| TD684 | $\Delta spoIVA::neo thrC::spoIVA^{T70A,T71A} \Delta spoVK::erm amyE::spoVK cat pyrD::spoVK-linker-GFP cat::tet$ | This study |
| TD836 | *ΔspoIVA::neo thrC::spoIVA<sup>K30E</sup> ΔspoVK::erm amyE::spoVK cat pyrD::spoVK-linker-GFP cat::tet* | This study |
| TD557 | *ΔspoIVA::neo thrC::spoIVA<sup>D97A</sup> spec amyE::spoVK cat* | This study |
| TD558 | *ΔspoIVA::neo thrC::spoIVA<sup>D97A</sup> spec amyE::spoVK<sup>A5V</sup> cat* | This study |
| TD563 | *ΔspoIVA::neo thrC::spoIVA<sup>T70A,T71A</sup> spec amyE::spoVK cat* | This study |
| TD564 | *ΔspoIVA::neo thrC::spoIVA<sup>T70A,T71A</sup> spec amyE::spoVK<sup>A5V</sup> cat* | This study |
| TD859 | *ΔspoIVA::neo thrC::spoIVA<sup>K30E</sup> spec amyE::spoVK cat* | This study |
| TD860 | *ΔspoIVA::neo thrC::spoIVA<sup>K30E</sup> spec amyE::spoVK<sup>A5V</sup> cat* | This study |
| TD574 | *ΔspoVK::erm amyE::spoVK<sup>Δ2-42</sup> cat* | This study |
| TD575 | *ΔspoVK::erm amyE::spoVK<sup>K105A,T106A</sup> cat* | This study |
| TD576 | *ΔspoVK::erm amyE::spoVK<sup>D162A,E163A</sup> cat* | This study |
| TD578 | *ΔspoVK::erm amyE::spoVK<sup>R218A,R276A</sup> cat* | This study |
| TD597 | *amyE::spoVK cat* | This study |
| TD598 | *amyE::spoVK<sup>Δ2-42</sup> cat* | This study |
| TD599 | *amyE::spoVK<sup>K105A,T106A</sup> cat* | This study |
| TD600 | *amyE::spoVK<sup>D162A,E163A</sup> cat* | This study |
| TD602 | *amyE::spoVK<sup>R218A,R276A</sup> cat* | This study |
| TD883 | *amyE::P<sub>spoVID</sub>-spoVK<sup>D162A,E163A</sup>-His<sub>6</sub> cat* | This study |
| TD884 | *amyE::P<sub>spoVID</sub>-spoVK<sup>D162A,E163A</sup>-FLAG cat* | This study |
| TD1193 | *amyE::P<sub>spoVID</sub>-spoVK<sup>A5V,D162A,E163A</sup>-FLAG cat* | This study |
| TD651 | *ΔspoVID::neo amyE::spoVK-linker-GFP cat* | This study |
| TD695 | *ΔspoVID::neo amyE::spoVID<sup>T532G</sup> cat pyrD::spoVK-linker-GFP cat::tet* | This study |
| TD1257 | *ΔspoIVA::erm ΔspoVID::neo thrC::spoIVA spec amyE::spoVID<sup>501-575</sup> pyrD::spoVK-linker-GFP cat::tet* | This study |
| KR394 | *ΔspoIVA::neo thrC::spoIVA spec* | Ramamurthi et al., 2006 |
| JB171 | *ΔspoIVA::erm ΔspoVID::neo thrC::spoIVA spec* | This study |
| JB174 | *ΔspoIVA::erm ΔspoVID::neo thrC::spoIVA spec amyE::spoVID cat* | This study |
| TD892 | *ΔspoIVA::erm ΔspoVID::neo thrC::spoIVA spec amyE::spoVID<sup>501-575</sup>* | This study |
| TD893 | *ΔspoIVA::erm ΔspoVID::neo thrC::spoIVA spec amyE::spoVID<sup>526-575</sup>* | This study |
| TD1268 | *ΔspoIVA::erm ΔspoVID::neo thrC::spoIVA spec amyE::P<sub>spoVID</sub>-spoVK<sup>D162A-E163A</sup>-FLAG cat* | This study |
| TD1281 | *ΔspoIVA::erm ΔspoVID::neo thrC::spoIVA spec amyE::P<sub>spoVID</sub>-spoVK<sup>D162A-E163A</sup>-FLAG cat sacA::GFP-spoVID<sup>501-575</sup> cat::tet* | This study |
| TD1267 | *ΔspoIVA::neo thrC::spoIVA spec amyE::P<sub>spoVID</sub>-spoVK<sup>D162A-E163A</sup>-FLAG cat* | This study |
| TD1277 | *ΔspoIVA::neo thrC::spoIVA spec amyE::P<sub>spoVID</sub>-spoVK<sup>D162A-E163A,Δ2-6</sup>-FLAG cat* | This study |
| TD1283 | *ΔspoIVA::neo thrC::spoIVA spec amyE::P<sub>spoVID</sub>-spoVK<sup>D162A-E163A,A5V</sup>-FLAG cat* | This study |
| TD1278 | *ΔspoIVA::erm ΔspoVID::neo thrC::spoIVA spec amyE::P<sub>spoVID</sub>-spoVK<sup>D162A-E163A,Δ2-6</sup>-FLAG cat* | This study |
| TD1284 | *ΔspoIVA::erm ΔspoVID::neo thrC::spoIVA spec amyE::P<sub>spoVID</sub>-spoVK<sup>D162A-E163A,A5V</sup>-FLAG cat* | This study |
| SC765 | *amyE::P<sub>spoVM</sub>-SpoVM-lphluorin-cat* | This study |
| SC766 | *amyE::P<sub>spoIIQ</sub>-lphluorin-cat* | This study |
| SC767 | *amyE::P<sub>spoVM</sub>-lphluorin-cat* | This study |
| SC777 | *thrC::P<sub>hyperspank</sub>-lphluorin-erm* | This study |

| *Escherichia coli* strains used in this study |  |  |
| --- | --- | --- |
| Strain | Genotype and description | Reference |
| BL21(DE3) | *pTD211* (*pET28a* backbone, *P<sub>T7</sub>-spoVK-His<sub>6</sub>*) | This study |
| BL21(DE3) | *pTD308* (*pET28a* backbone, *P<sub>T7</sub>-spoVK<sup>D162A,E163A</sup>-His<sub>6</sub>*) | This study |

**Figure S1.**
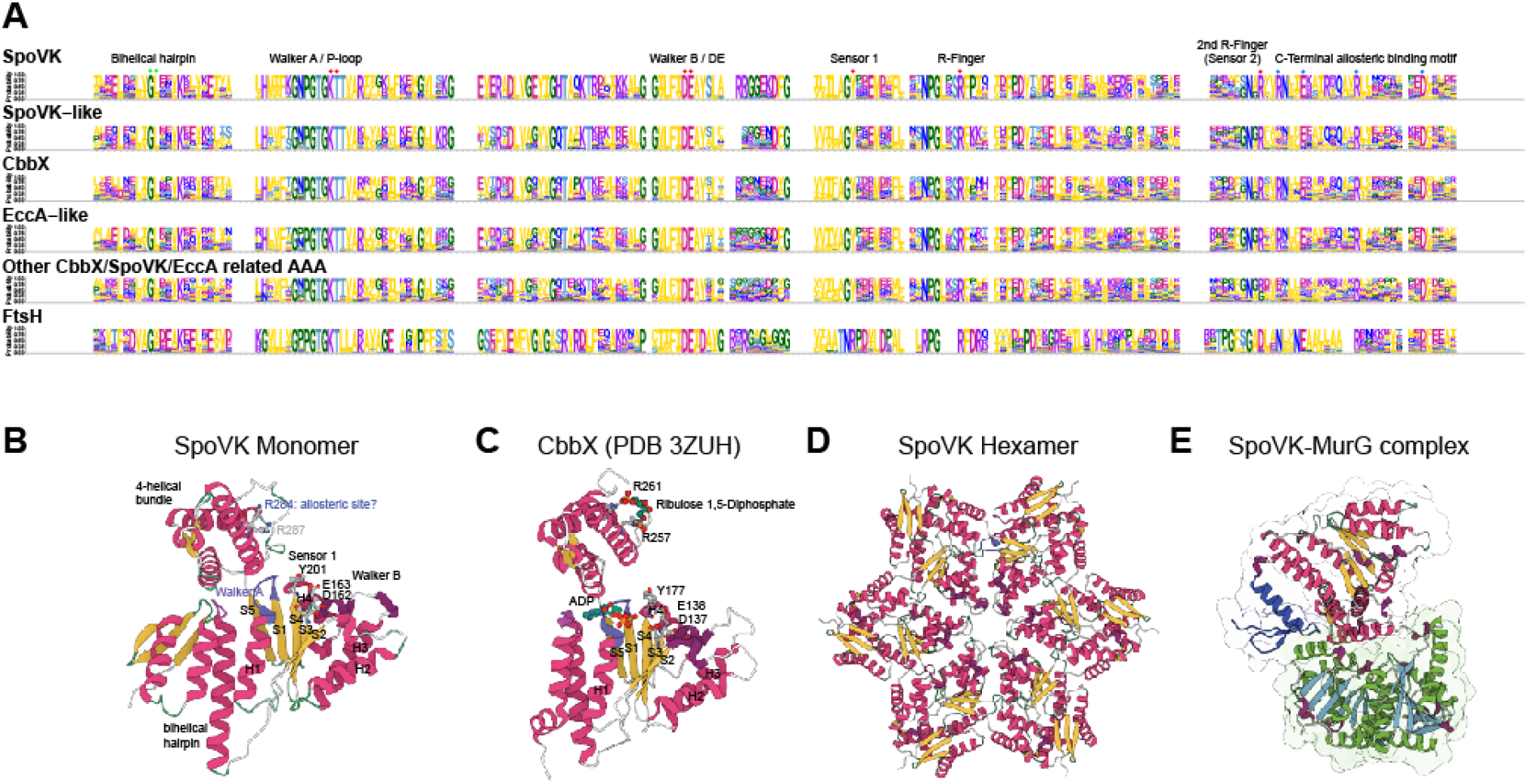
Sequence and structural comparison between SpoVK and members of the SpoVK-EccA-like AAA+ ATPase clade. (A) Sequence logo displaying conservation of amino acid residues in the SpoVK-EccA-like AAA+ ATPase clade. Letters represent amino acid abbreviations; height of each letter represents the probability of conservation among orthologs of the family. The first and second arginine fingers are marked with red dots. Blue dots are conserved residues contacting the allosteric ligand ribulose bisphosphate. (B-E) Cartoon depictions of (B) AlphaFold2 prediction of SpoVK structure, (C) CbbX from *Cereibacter sphaeroides* (PDB 3ZUH), (D) AlphaFold-Multimer prediction of SpoVK hexamer, and (E) AlphaFold-Multimer prediction of SpoVK-MurG complex.

**Figure S2.**
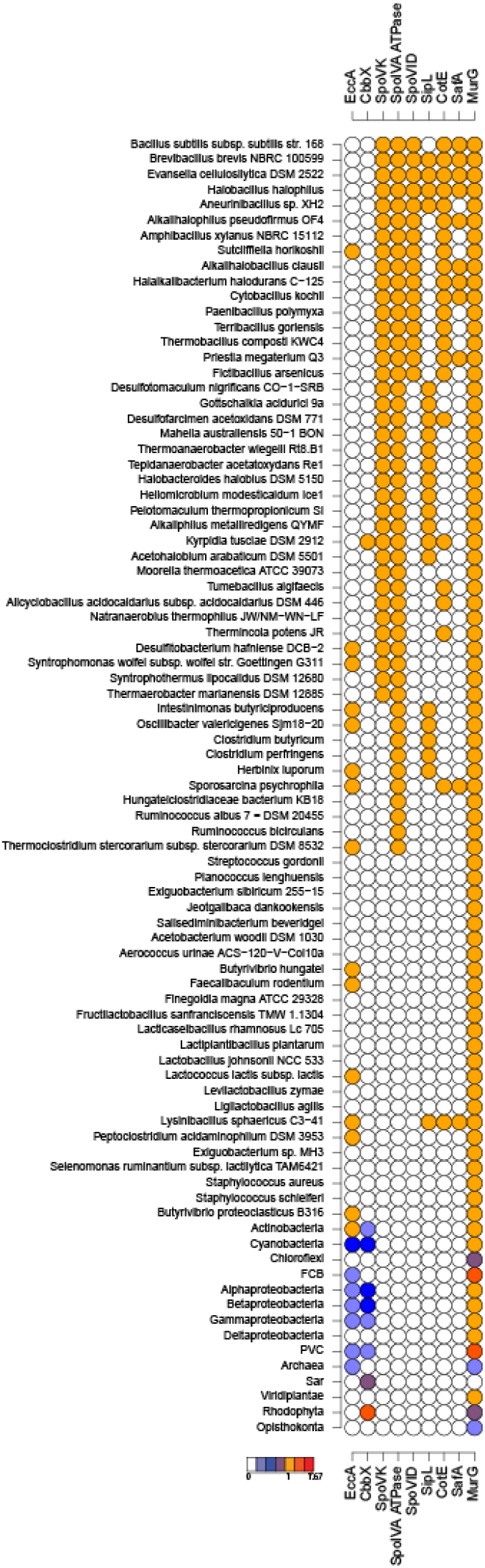
Phyletic pattern vectors of EccA, CbbX, SpoVK, SpoIVA ATPase, SpoVID, SipL, CotE, SafA, and MurG domain proteins. The presence or absence of the protein in the clade or organism (Firmicutes) are shown. The legend shows the color gradient for normalized counts that were scaled by square root.

**Figure S3.**
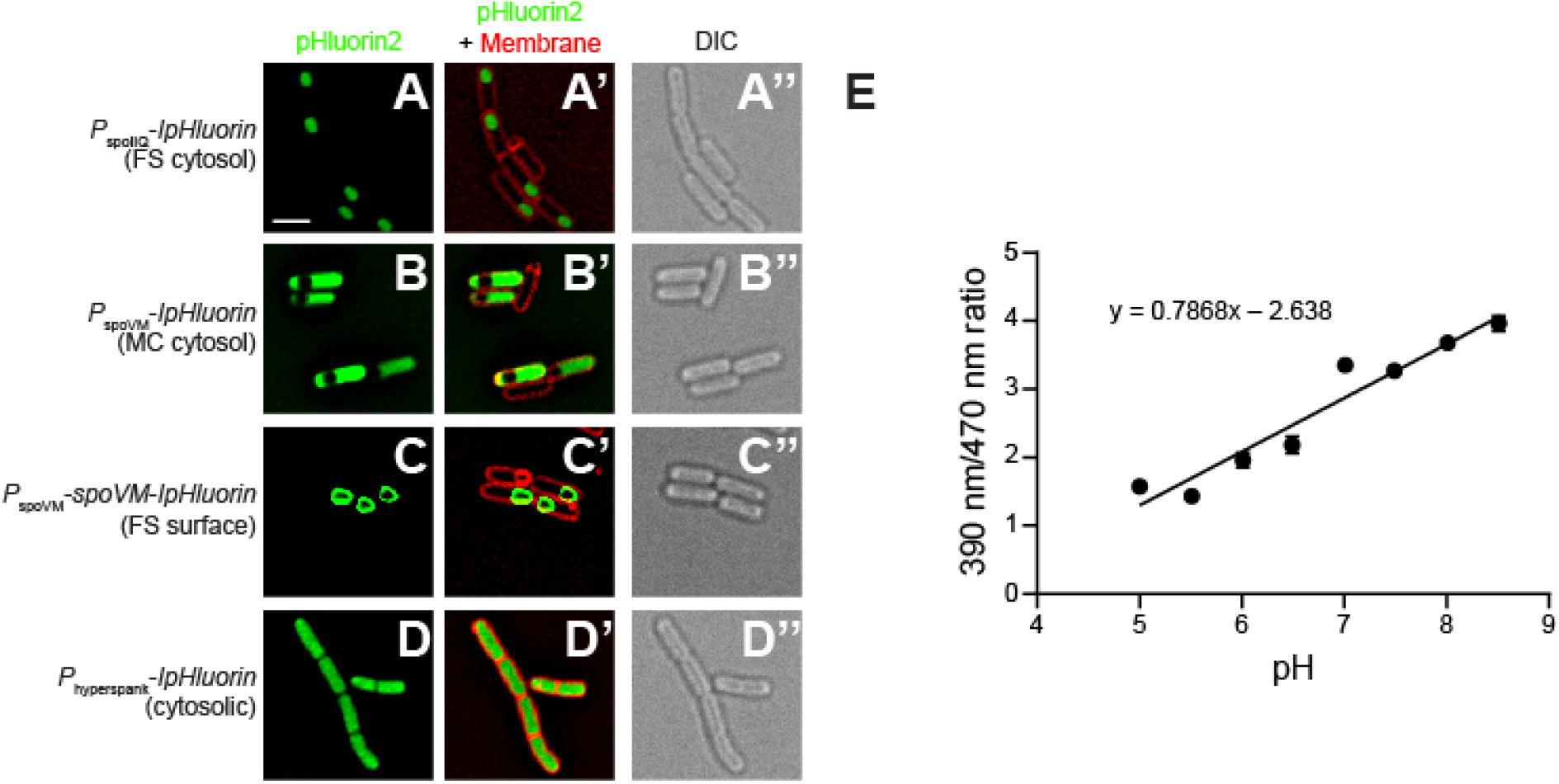
Subcellular localization of various IpHluorin constructs during sporulation. Fluorescence micrographs of *B. subtilis* (A-C’’) at t = 4 h after induction of sporulation producing (A-A’’) IpHluorin in the forespore, expressed under control of the *spoIIQ* promoter, (B-B’’) IpHluorin in the mother cell, expressed under control of the *spoVM* promoter, or (C-C’’) SpoVM-IpHluorin at the forespore surface, expressed in the mother cell under control of the *spoVM* promoter; or (D-D’’) during vegetative growth producing IpHluorin, expressed under control of an IPTG-inducible promoter at t = 2 h after IPTG induction. (A-D) fluorescence from IpHluorin; (A’-D’) overlay, IpHluorin and membranes visualized using FM4-64; (A’’-D’’) differential interference contrast. Strains: SC765, SC766, SC767, and SC777. Scale bar: 2 µm. (E) Calibration curve of ratio of fluorescence emission at 510 nm when excited at either 390 nm or 470 nm ratio as a function of media pH.

## Notes

### Competing Interest Statement

The authors have declared no competing interest.

